# The deubiquitinase Ubp3/Usp10 constrains glucose-mediated mitochondrial repression via phosphate budgeting

**DOI:** 10.1101/2022.12.29.522272

**Authors:** Vineeth Vengayil, Shreyas Niphadkar, Swagata Adhikary, Sriram Varahan, Sunil Laxman

## Abstract

Many cells in high glucose repress mitochondrial respiration, as observed in the Crabtree and Warburg effects. Our understanding of biochemical constraints for mitochondrial activation is limited. Using a *Saccharomyces cerevisiae* screen, we identified the conserved deubiquitinase Ubp3 (Usp10), as necessary for mitochondrial repression. Ubp3 mutants have increased mitochondrial activity despite abundant glucose, along with decreased glycolytic enzymes, and a rewired glucose metabolic network with increased trehalose production. Utilizing *Δubp3* cells, along with orthogonal approaches, we establish that the high glycolytic flux in glucose continuously consumes free Pi. This restricts mitochondrial access to inorganic phosphate (Pi), and prevents mitochondrial activation. Contrastingly, rewired glucose metabolism with enhanced trehalose production and reduced GAPDH (as in *Δubp3* cells) restores Pi. This collectively results in increased mitochondrial Pi and derepression, while restricting mitochondrial Pi transport prevents activation. We therefore suggest that glycolytic-flux dependent intracellular Pi budgeting is a key constraint for mitochondrial repression.

## Introduction

Rapidly proliferating cells have substantial metabolic and energy demands in order to increase biomass (Cai and Tu, 2012; Zhu and Thompson, 2019). This includes a high ATP demand, obtained from cytosolic glycolysis or mitochondrial oxidative phosphorylation (OXPHOS), to fuel multiple reactions (Nelson et al., 2008). Interestingly, many rapidly proliferating cells preferentially rely on ATP from glycolysis/fermentation over mitochondrial respiration even in oxygen-replete conditions, and is the well-known Warburg effect (Vander Heiden et al., 2009; Warburg, 1925). Many such cells repress mitochondrial processes in high glucose, termed glucose-mediated mitochondrial repression or the Crabtree effect (Crabtree, 1929; De Deken, 1966). This is observed in some tumors (Vander Heiden et al., 2009), neutrophils (Xia et al., 2021), activated macrophages (Kornberg, 2020), stem cells (Abdel-Haleem et al., 2017; Pacini and Borziani, 2014; Tsogtbaatar et al., 2020), and famously *Saccharomyces cerevisiae* (De Deken, 1966). Numerous studies have identified signaling programs or regulators of glucose-mediated mitochondrial repression. However, biochemical programs and regulatory processes in biology evolve around key biochemical constraints (Cornish-Bowden, 2016). The biochemical constraints for mitochondrial repression remain unresolved (Diaz-Ruiz et al., 2011; Hammad et al., 2016).

There are two hypotheses on the biochemical principles driving mitochondrial repression. The first proposes direct roles for glycolytic intermediates in driving mitochondrial respiration, by inhibiting specific mitochondrial outputs (Díaz-Ruiz et al., 2008; Lemus et al., 2018). The second hypothesizes that a competition between glycolytic and mitochondrial processes for mutually required metabolites/co-factors such as pyruvate, ADP or inorganic phosphate (Pi) could determine the extent of mitochondrial repression (Diaz-Ruiz et al., 2011; Hammad et al., 2016; Koobs, 1972). These are not all mutually exclusive, and a combination of these factors might dictate mitochondrial repression in high glucose. However, any hierarchies of importance are unclear (Rodríguez-Enríquez et al., 2001), and experimental data for the necessary constraints for mitochondrial repression remains incomplete.

One approach to resolve this question has been to identify regulators of metabolic state under high glucose. Post-translational modifications (PTMs) regulated by signaling systems can regulate mitochondrial repression (Broach, 2012; Hitosugi and Chen, 2014; Tripodi et al., 2015). Ubiquitination is a PTM that regulates global proteostasis (Hershko and Ciechanover, 1998; Komander and Rape, 2012), but the roles of ubiquitination-dependent processes in regulating mitochondrial repression are poorly explored. Ubiquitination itself is determined by the balance between ubiquitination, and deubiquitinase (DUB) dependent deubiquitination (Pickart and Eddins, 2004). Little is known about the roles of DUBs in regulating metabolic states, making the DUBs interesting candidate regulators of mitochondrial repression.

In this study, using an *S. cerevisiae* DUB knock-out library-based screen, we identified the evolutionarily conserved deubiquitinase Ubp3 (mammalian Usp10) as required for mitochondrial repression in high glucose. Loss of Ubp3 resulted in mitochondrial activation, along with a reduction in the glycolytic enzymes-phosphofructokinase 1 (Pfk1) and GAPDH (Tdh2 and Tdh3). This consequently reroutes glucose flux and increases trehalose biosynthesis. This metabolic rewiring increases Pi release from trehalose synthesis, and decreases Pi consumption in glycolysis, to cumulatively increase Pi pools. Using *ubp3Δ* cells along with independent analysis of wild-type cells, and isolated mitochondrial fractions, we establish that glycolytic-flux dependent Pi allocations to mitochondria determines mitochondrial activity. Through these data, we propose how intracellular Pi balance as controlled by glycolytic flux, is a biochemical constraint for mitochondrial repression.

## Results

### A deubiquitinase deletion screen identifies Ubp3 as a regulator of glucose-mediated mitochondrial repression

In cells such as *S. cerevisiae*, high glucose represses mitochondrial activity as well as OXPHOS-dependent ATP synthesis (De Deken, 1966; Postma et al., 1989) (illustrated in Figure 1A). Our initial objective was to identify proteostasis regulators of glucose-mediated mitochondrial repression. We generated and used a deubiquitinase (DUB) deletion strain library of *S. cerevisiae* (Figure 1-figure supplement 1A), to unbiasedly identify regulators of mitochondrial repression by measuring the fluorescence intensity of a potentiometric dye Mitotracker CMXRos (illustrated in Figure 1B). Using this screen, we identified DUB mutants with altered mitochondrial membrane potential (Figure 1C, Figure 1-figure supplement 1 A). Note: WT cells in a respiratory medium (2% ethanol) were used as a control to estimate maximum mitotracker fluorescence intensity (Figure 1-figure supplement 1B).

**Figure 1.**
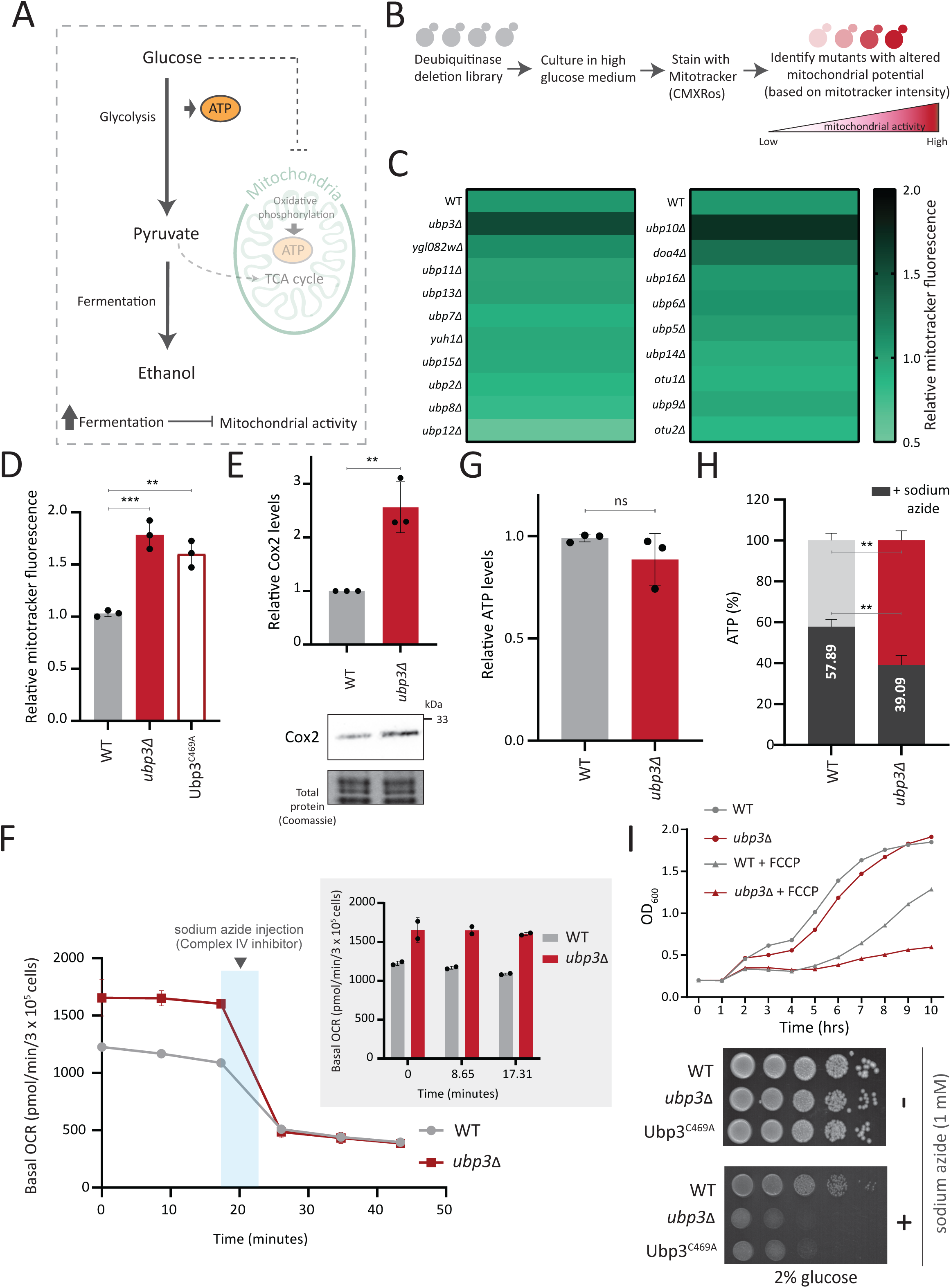
A deubiquitinase deletion screen identifies Ubp3 as a regulator of glucose-mediated mitochondrial repression. A) Schematic depicting glucose-mediated mitochondrial repression (Crabtree effect). B) Schematic describing the screen with a yeast DUB KO library to identify regulators of Crabtree effect. C) Identifying DUB knockouts with altered mitochondrial potential. Heat map shows relative mitochondrial membrane potential of 19 deubiquitinase deletions in high glucose, from 2 biological replicates. Also see Figure 1-figure supplements 1A, B. D) The deubiquitinase activity of Ubp3 and repression of mitochondrial membrane potential. WT, *ubp3Δ,* and Ubp3^C469A^ were grown inhigh glucose and relative mitochondrial membrane potential was measured. Data represent mean ± SD from three biological replicates (n=3). Also see Figure 1-figure supplement 1 D. E) Effect of loss of Ubp3 on ETC complex IV subunit Cox2. WT and *ubp3Δ* were grown in high glucose, and Cox2 was measured (western blot using an anti-Cox2 antibody). A representative blot (out of 3 biological replicates, n=3) and their quantifications are shown. Data represent mean ± SD. F) Basal oxygen consumption rate (OCR) in high glucose in *ubp3Δ*. WT and *ubp3Δ* were grown in high glucose, and OCR was measured. Basal OCR corresponding to ∼3 × 10^5 cells, from two independent experiments (n=2), normalized to the OD600 is shown. Bar graph representations are shown in the inset. Data represent mean ± SD. G) Total ATP levels in *ubp3Δ* and WT. WT and *ubp3Δ* were grown in high glucose, and total ATP were measured. Data represent mean ± SD from three biological replicates (n=3). H) Dependence of *ubp3Δ* on mitochondrial ATP. WT and *ubp3Δ* cells were grown in high glucose, and treated with 1 mM sodium azide for 45 minutes. Total ATP levels in sodium azide treated and untreated cells were measured. Data represent mean ± SD (n=3). I) Requirement for mitochondrial respiration in high glucose in *ubp3Δ*. A growth curve of WT and *ubp3Δ* in high glucose in the presence of OXPHOS uncoupler FCCP (10 µM), and serial dilution growth assay in high glucose in the presence/absence of sodium azide (1 mM) are shown. Data represent mean ± SD (n=2). Also see Figure 1-figure supplement 1 H, I. Data information: **p<0.01, ***p<0.001.

A prominent ‘hit’ was the evolutionarily conserved deubiquitinase Ubp3 (Figure 1C), (homologous to mammalian Usp10) (Figure 1-figure supplement 1D). Due to its high degree of conservation across eukaryotes (Figure 1-figure supplement 1D) as well as putative roles in metabolism or mitochondrial function (Isasa et al., 2015; Nostramo et al., 2016; Ossareh-Nazari et al., 2010), we focused our further attention on this DUB. Cells lacking Ubp3 showed an ∼1.5-fold increase in mitotracker fluorescence (Figure 1C, Figure 1-figure supplement 1A, C). Cells with catalyticaly inactive Ubp3 (Ubp3^C469A^), showed increased mitochondrial potential comparable to *ubp3Δ* (Figure 1D). This data confirmed that Ubp3 catalytic activity of is required to fully repress mitochondrial activity under high glucose. The catalytic site mutation did not affect steady-state Ubp3 levels (Figure 1-figure supplement 1E).

Next, to assess the requirement of Ubp3 for mitochondrial function, we quantified the electron transport chain (ETC) complex IV subunit Cox2 (Fontanesi et al., 2006)(Figure 1E). *ubp3Δ* had higher Cox2 than WT (Figure 1E). As a control, we estimated total mitochondrial content in WT and *ubp3Δ* cells, using either estimates of the structural protein Tom70, or measuring the fluorescence-intensity in strains engineered with mitochondrial targeted mNeonGreen (Dua et al., 2022). There was no increase in the total mitochondrial volume (estimated by measuring the intensity of mitochondria targeted mNeon green) (Figure 1-figure supplement 1F) or mitochondrial outer membrane protein Tom70 (Figure 1-figure supplement 1G). This suggests that the increased Cox2 is not merely because of higher total mitochondrial content. We next measured the basal oxygen consumption rate (OCR) of *ubp3Δ*, and basal OCR was higher in *ubp3Δ*, indicating higher respiration (Figure 1F).

Next, we asked if mitochondrial ATP synthesis was higher in *ubp3Δ*. The total ATP levels in WT and *ubp3Δ* were comparable (Figure 1G). However, upon treatment with the ETC complex IV inhibitor sodium azide, ATP levels in WT were higher than *ubp3Δ*, contributing to ∼60% of the total ATP (Figure 1H). In contrast, the ATP levels in *ubp3Δ* after sodium azide treatment were ∼ 40% of the total ATP (Figure 1H). These data suggest a higher contribution of mitochondrial ATP synthesis towards the total ATP pool in *ubp3Δ*.

We next asked if *ubp3Δ* required higher mitochondrial activity for growth, using a series of mitochondrial activity inhibitors and comparing relative growth. In high glucose, WT cells show minimal growth inhibition in the presence of sodium azide, indicating lower reliance on mitochondrial function (Figure 1I). Contrastingly, *ubp3Δ* or Ubp3^C469A^ show a severe growth defect in the presence of the mitochondrial ETC complex inhbitors sodium azide and oligomycin, and the mitochondrial OXPHOS uncoupler FCCP (Figure 1I, Figure 1-figure supplement 1H). Additionally, the loss of Ubp3 in respiration defective cox2-62 cells (Bonnefoy et al., 2001), or Rho0 cells (which lacks mitochondrial DNA) resulted in a severe growth defect (Figure 1-figure supplement 1I). Deletion of ATP synthase subunits Atp1 and Atp10 also results in a severe growth defect in *ubp3Δ* compared to WT (Figure 1-figure supplement 1I). Together, these results indicate that the loss of Ubp3 makes cells dependent on mitochondrial ATP synthesis in high glucose.

In order to address whether the deletion of Ubp3 might increase ubiquitinated proteins and consequent proteostatic stress, we analysed the global ubquitination state in *ubp3Δ* cells. WT and *ubp3Δ* cells grown under brief (1 hour) heat stress, which increases protein ubiquitination, was used as a control. However, we did not observe a significant increase in the global ubiqutinatin state in *ubp3Δ* cells (Figure 1-figure supplement 1J). This suggests that the altered mitochondrial metabolism in *ubp3Δ* cells is unlikely to be due to proteostatic stress.

Collectively these data show that in 2% glucose, *ubp3Δ* have high mitochondrial activity, respiration, and rely on this mitochondrial function for ATP production and growth. We therefore decided to use *ubp3Δ* cells to start delineating requirements for glucose-mediated mitochondrial repression.

### Key glycolytic enzymes decrease and glucose flux is rerouted in *ubp3Δ* cells

Glucose-6 phosphate (G6P) is the central node in glucose metabolism where carbon-allocations are made towards distinct metabolic arms, primarily: glycolysis, the pentose phosphate pathway (PPP), and trehalose biosynthesis (Figure 2A). We first compared amounts of two key (‘rate-controling’) glycolytic enzymes - Phosphofructokinase 1 (Pfk1), GAPDH isozymes (Tdh2, Tdh3) (Nelson et al., 2008), along with the enolase isozymes (Eno1, Eno2) in WT and *ubp3Δ* cells. Pfk1, Tdh2, and Tdh3 substantially decreased in *ubp3Δ* (but not Eno1 and Eno2) (Figure 2B, Figure 2-figure supplement 1A). Since deubiquitinases can control protein amounts by regulating proteasomal degradation, we asked if the decrease in Pfk1, Tdh2 and Tdh3 in *ubp3Δ* is due to proteasomal degradation. To test this, we measured the levels of these enzymes in *ubp3Δ* after treatement with proteasomal inhibitor MG132. We did not observe any rescue in the levels of these enzymes in MG132 treated samples, suggesting that the decreased levels of these enzymes were not due to increased proteasomal degradation (Figure 2-figure supplement 1B). To further understand if these enzyme transcripts are altered in *ubp3Δ*, we measured the mRNA levels of PFK1, TDH2 and TDH3 in WT, *ubp3Δ* and Ubp3^C469A^ cells. The transcripts of all the three genes in *ubp3Δ* and Ubp3^C469A^ cells decreased (Figure 2-figure supplement 1C), suggesting that Ubp3 regulates the transcripts of these glycolytic enzyme genes. The reduction in the Pfk and GAPDH enzyme amounts was intriguing, because the Pfk and GAPDH steps are critical in determining glycolytic flux (Nelson et al., 2008; Shestov et al., 2014). A reduction in these enzymes could therefore decrease glycolytic flux, and would reroute glucose (G6P) allocations via mass action towards other branches of glucose metabolism, primarily the pentose phosphate pathway as well as trehalose biosynthesis (Figure 2A). To assess this, we first measured the steady-state levels of key glycolytic and pentose phosphate pathway (PPP) intermediates, and trehalose in WT or *ubp3Δ* using targeted LC-MS/MS (Figure 2C, Figure 2-figure supplement 1D). Glucose-6/fructose-6 phosphate (G6P/F6P) increased in *ubp3Δ* (Figure 2C). Concurrently, trehalose, and the PPP intermediates ribose 5-phosphate (R5P) and sedoheptulose 7-phosphate (S7P), increased in *ubp3Δ* (Figure 2C).

**Figure 2:**
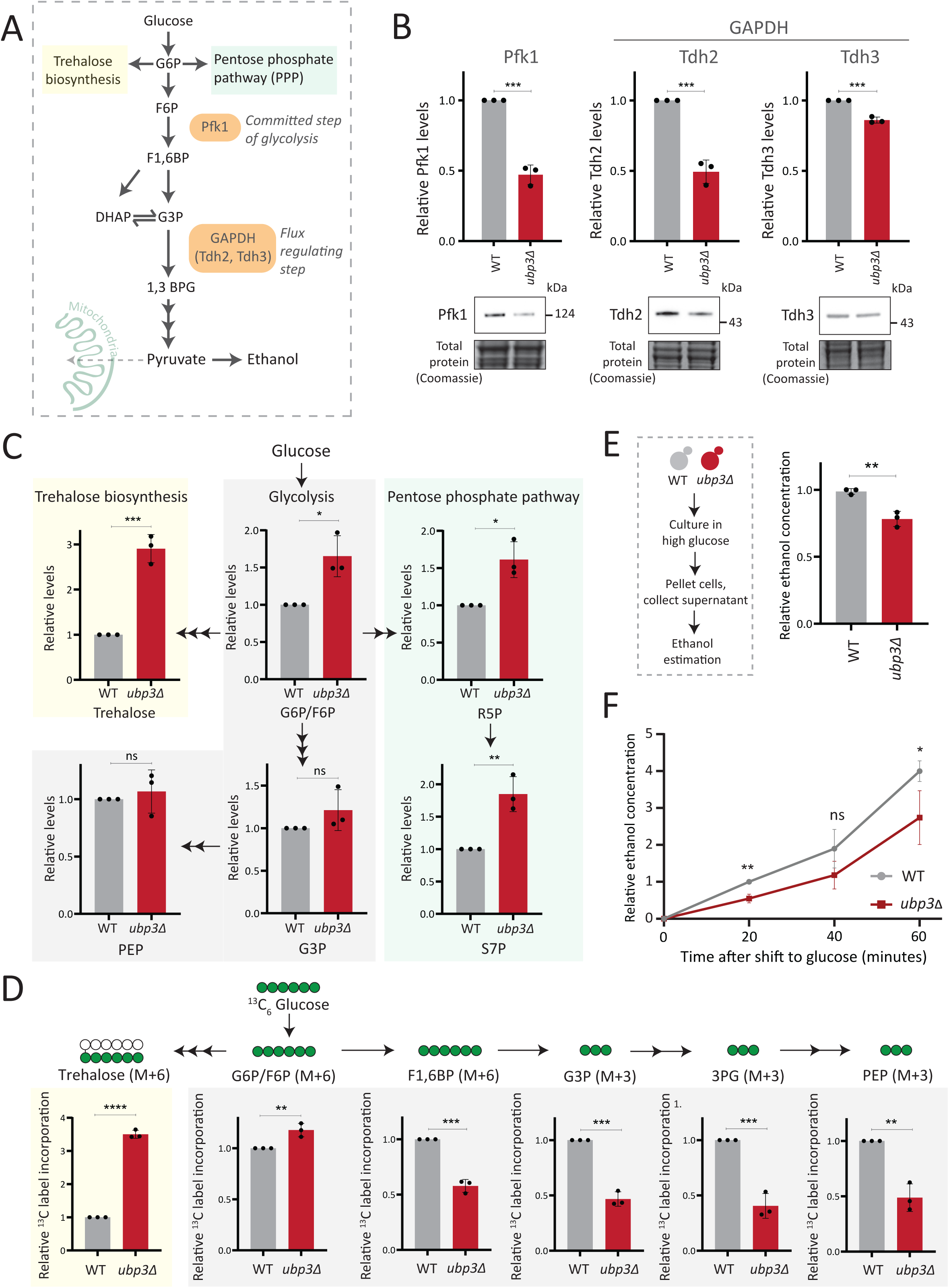
Key glycolytic enzymes decrease and glucose flux is rerouted in *ubp3Δ* cells. A) A schematic illustrating directions of glucose-6-phosphate (G6) flux in cells. Glucose is converted to G6P, a precursor for trehalose, the pentose phosphate pathway (PPP), and glycolysis. B) Effect of loss of Ubp3 on key glycolytic enzymes. WT and *ubp3Δ* were grown in high glucose and the Pfk1, Tdh2, and Tdh3 levels were measured by western blot using an anti-FLAG antibody. A representative blot (out of three biological replicates, n=3) and their quantification are shown. Data represent mean ± SD. Also see Figure 2-figure supplement 1A. C) Steady-state metabolite amounts in WT and *ubp3Δ* in high glucose. Relative steady-state levels of trehalose, major glycolytic, and PPP intermediates were estimated in WT and *ubp3Δ*. Data represent mean ± SD from three biological replicates (n=3). Also see Appendix Table S3. D) Relative glycolytic and trehalose synthesis flux in WT and *ubp3Δ*. Relative ^13^C-label incorporation into trehalose and glycolytic intermediates, after a pulse of 1% ^13^C_6_ glucose is shown. Data represent mean ± SD from three biological replicates (n=3). Also see Appendix Table S3, Figure 2-figure supplement 1 D, E. E) Ethanol production in *ubp3Δ*. WT and *ubp3Δ* were grown in high glucose and ethanol in the media was measured. Data represent mean ± SD from three biological replicates (n=3). F) Relative rate of ethanol production in WT vs *ubp3Δ*. WT and *ubp3Δ* were grown in high glucose (to OD_600_ ∼ 0.6), equal numbers of cells were shifted to fresh medium (high glucose) and ethanol concentration in the medium was measured temporally. Data represent mean ± SD from three biological replicates (n=3) Data information: *p<0.05, **p<0.01, ***p<0.001.

Since steady-state metabolite amounts cannot separate production from utilization, in order to unambiguously assess if glycolytic flux is reduced in *ubp3Δ* cells, we utilized a pulse labeling of ^13^C_6_ glucose, following which the label incorporation into glycolytic and other intermediates was measured. Note that because glycolytic flux is very high in yeast, this experiment would require rapid pulsing and extraction of metabolites in order to stay in a linear range and avoid label saturation. We therefore established a very short time point of label-addition, quenching and metabolite extraction post ^13^C glucose pulse. Since flux saturates/reaches steady-state in seconds, we first ensured that the label incorporation into individual metabolites after the ^13^C glucose pulse was in the linear range, and for early glycolytic intermediates this was seconds after glucose addition. This new methodology is extensively described in materials and methods, with required controls shown in Figure 2-figure supplement 1F. WT and *ubp3Δ* cells were grown in high glucose, pulsed with ^13^C_6_ glucose, and the relative ^13^C label incorporation into glycolytic intermediates and trehalose were measured, as shown in the schematic (Figure 2-figure supplement 1E). In *ubp3Δ,* ^13^C label incorporation into G6P/F6P as well as trehalose substantially increased (Figure 2D). Contrastingly, ^13^C label incorporation into glycolytic intermediates F1,6BP, G3P, 3PG and PEP decreased, indicating decreased glycolytic flux (Figure 2D). We next measured ethanol concentrations and production rates, as an additional output of relative glycolytic rates. We observed decreased steady-state ethanol levels, as well as ethanol production rates in *ubp3Δ* (Figure 2F).

Glycolysis-derived pyruvate is transported to mitochondria and fuels the TCA cycle. Therefore, we asked if the decreased glycolytic rate result in a decrease in the TCA cycle flux as well. To test this, we first measured the steady state levels of TCA cycle intermediates in WT or *ubp3Δ* using targeted LC-MS/MS. We did not observe any significant change in the levels of TCA cycle intermediates in *ubp3Δ,* except malate which showed a significant decrease in *ubp3Δ* (Figure 2-figure supplement 2A). Next, in order to assess if TCA cycle flux reduces in *ubp3Δ* cells, WT and *ubp3Δ* cells were grown in high glucose, pulsed with ^13^C_6_ glucose, and the relative ^13^C label incorporation into TCA cycle intermediates was measured, as shown in the schematic (Figure 2-figure supplement 2B). The kinetics of ^13^C label incorporation in TCA cycle intermediates are shown in Figure 2-figure supplement 2C. We did not observe any significant change in the relative ^13^C label incorporation in TCA cycle intermediates in *ubp3Δ.* Therefore, these data suggest that the decreased glycolytic flux in *ubp3Δ* does not result in a decrease in TCA cycle flux. The increased respiration and mitochondrial activity in *ubp3Δ* cells is therefore driven via other factors.

Collectively, these results reveal that that reduced Pfk1 and GAPDH (in *ubp3Δ*) decrease glucose flux via glycolysis, which results in rewired glucose flux towards trehalose biosynthesis and the PPP.

### Rerouted glucose flux results in phosphate (Pi) accumulation

We therefore asked if the proteomic state, as observed in *ubp3Δ* cells, could provide clues to explain the coupling between mitochondrial derepression and rerouted glucose flux. A recent study by Isasa et al., had systematically quantified the changes in protein levels in *ubp3Δ* cells (Isasa et al., 2015). We therefore reanalyzed this extensive dataset, looking for changes in proteins that would correlate with these metabolic processes. Notably, we observed increased levels of proteins of the mitochondrial ETC and respiration, and decreased amounts of glucose metabolizing enzymes in *ubp3Δ* (Figure 3A). Additionally, multiple proteins involved in regulating phosphate (Pi) homeostasis were decreased in *ubp3Δ* (Figure 3A) (Isasa et al., 2015). We had recently uncovered a reciprocal coupling of Pi homeostasis with the different arms of glucose metabolism, particularly trehalose biosynthesis (Gupta et al., 2019; Gupta and Laxman, 2021), and therefore wondered if a glycolytic-flux dependent change in Pi homeostatsis had any role in mitochondrial respiration. Our hypothesis was refined based on the reasoning given below.

**Figure 3:**
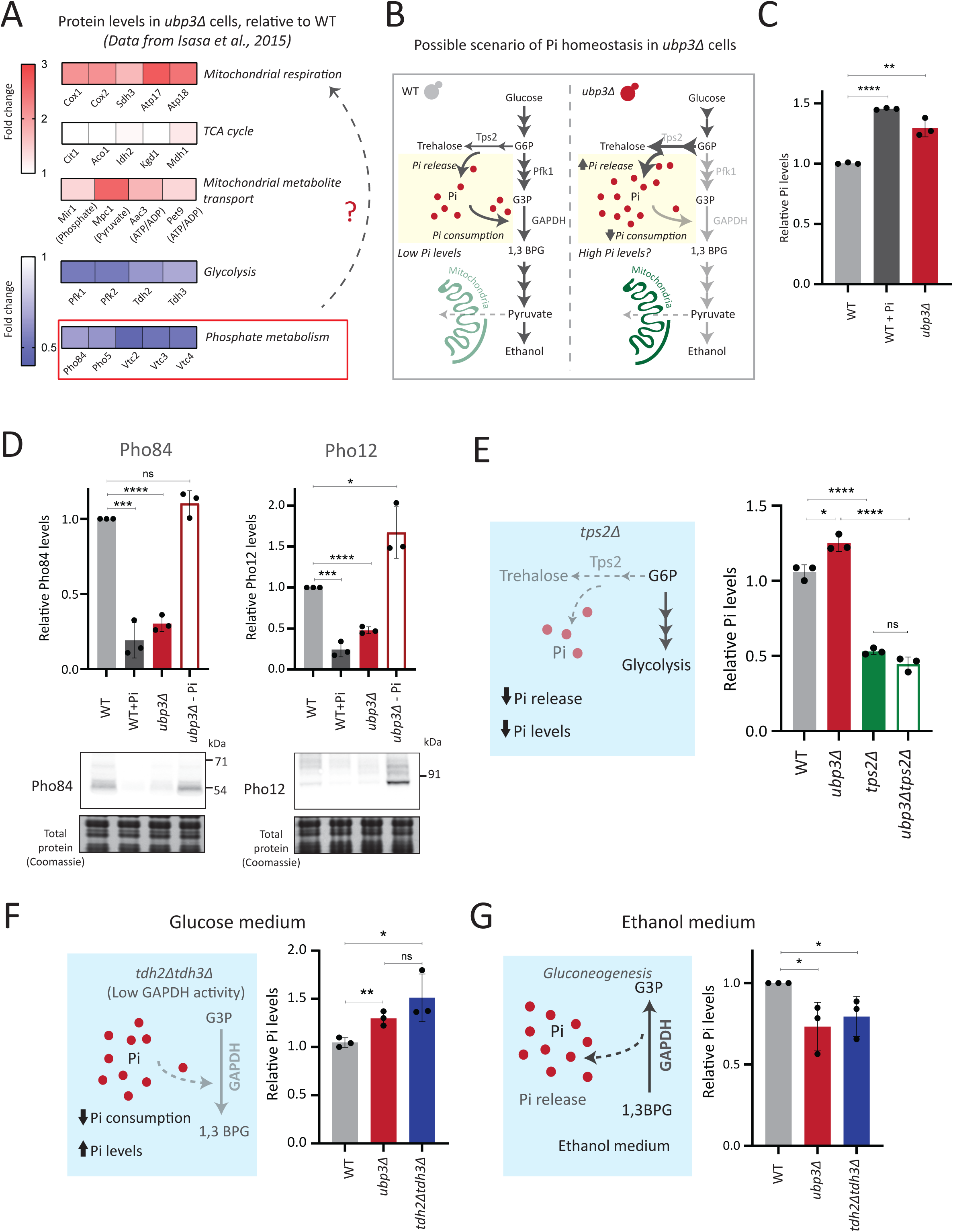
Rerouted glucose flux results in phosphate (Pi) accumulation. A) Changes in protein levels in *ubp3Δ* (dataset from Isasa et al., 2015)*. ubp3Δ* cells have an increase in proteins involved in mitochondrial respiration and decrease in proteins involved in glucose and phosphate metabolism. B) Schematic showing maintenance of Pi balance during glycolysis. Trehalose synthesis from G6P releases Pi, and the conversion of G3P to 1,3BPG by GAPDH consumes Pi. In *ubp3Δ*, trehalose biosynthesis (which releases Pi) increases. *ubp3Δ* have decreased GAPDH, which will decrease Pi consumption. This increase in Pi release along with decreased Pi consumption could increase cytosolic Pi. C) Intracellular Pi levels in WT and *ubp3Δ*. WT and *ubp3Δ* were grown in high glucose and the total free phosphate (Pi) levels were estimated. WT in high Pi (2% glucose, 10mM Pi) was a positive control. Data represent mean ± SD from three biological replicates (n=3). Also see Figure 3-figure supplement 1A. D) Pho regulon responses in WT and *ubp3Δ*. Protein levels of Pho84-FLAG and Pho12-FLAG were compared between WT grown in high glucose and in high Pi, *ubp3Δ* in high glucose with or without a shift to a no-Pi medium for one hour, by western blot. A representative blot (out of three biological replicates, n=3) and their quantifications are shown. Data represent mean ± SD. E) Contribution of trehalose synthesis as a Pi source. WT, *tps2Δ*, *ubp3Δ,* and *ubp3Δtps2Δ* were grown in high glucose and the total Pi levels were estimated. Data represent mean ± SD from three biological replicates (n=3). Also see Figure 3-figure supplement 1B. F) Loss of GAPDH isozymes Tdh2 and Tdh3 and effect on Pi. WT, *ubp3Δ,* and *tdh2Δtdh3Δ* were grown in high glucose and total Pi was estimated. Data represent mean ± SD from three biological replicates (n=3). G) Pi levels in *ubp3Δ* and *tdh2Δtdh3Δ* cells in ethanol medium. WT, *ubp3Δ,* and *tdh2Δtdh3Δ* cells were grown in ethanol medium and the total Pi levels were estimated. Data represent mean ± SD from three biological replicates (n=3). Data information: *p<0.05, **p<0.01, ***p<0.001, ****p<0.0001.

One explanation for mitochondrial repression can be an internal competition for shared metabolites/co-factors between (cytosolic) glycolytic and mitochondrial processes, that mitochondria might not be sufficiently able to access (Diaz-Ruiz et al., 2011; Koobs, 1972; Rodríguez-Enríquez et al., 2001). In this context, a plausible role for inorganic phosphate (Pi) in regulating mitochondrial repression can be hypothesized. Cytosolic glycolysis requires rapid, high consumption of net Pi (Mason et al., 1981; Rodríguez-Enríquez et al., 2001; Van Heerden et al., 2014), and this could possibly limit the Pi that is continuously available for mitochondrial use (Brazy et al., 1982; Koobs, 1972), thereby repressing mitochondria. Contextually, the balance between reactions releasing vs consuming Pi could explain changes in global Pi levels (Gupta and Laxman, 2021). A continuous hub of Pi consumption is glycolysis. Here, GAPDH catalyzes G3P to 1,3BPG, converting ADP to ATP, while concurrently consuming a molecule of Pi (Hohmann et al., 1996; Van Heerden et al., 2014). This Pi that goes into ATP will subsequently be used for nucleotide biosynthesis, polyphosphate biosynthesis and protein phosphorylation (Gupta and Laxman, 2021; Hunter, 2012). Therefore, we can surmise that in high glycolytic flux, the production of ATP, nucleotides and polyphosphates is concurrent with Pi consumption (Austin and Mayer, 2020; Hohmann et al., 1996; Ljungdahl and Daignan-Fornier, 2012). Could this reaction therefore limit cytosolic Pi for the mitochondria (as illustrated in Figure 3B)? Notably, *ubp3Δ* have reduced GAPDH levels and decreased glycolytic flux. Second, trehalose synthesis is a Pi releasing reaction, and a major source of free Pi that is critical for Pi homeostasis (Gupta et al., 2019; Van Heerden et al., 2014). Flux through this reaction is also substantially higher in *ubp3Δ*. Therefore, these cells might have increased Pi release (via trehalose), coupled with decreased Pi consumption (via decreased GAPDH). We therefore asked if total Pi increases in *ubp3Δ* (Figure 3B)?

To test this, we directly assessed total Pi levels in *ubp3Δ* and WT. *ubp3Δ* cells had higher Pi than WT, and this was comparable to Pi in WT grown in excess Pi (Figure 3C). Similarly, Pi amounts also increased in Ubp3^C469A^ (Figure 3-figure supplement 1A). Therefore, the loss of Ubp3 increases intracellular Pi levels. Next, we asked if *ubp3Δ* cells exhibit signatures of a ‘high Pi’ state. *S. cerevisiae* maintains internal Pi balance by controlling the expression of multiple genes collectively known as the Pho regulon (Mouillon and Persson, 2006). The Pho regulon is induced under Pi limitation, and repressed during Pi sufficiency (Gupta et al., 2019; Mouillon and Persson, 2006). We assessed two major Pho proteins (Pho84: a high-affinity membrane Pi transporter, and Pho12: an acid phosphatase) in WT and *ubp3Δ* in high glucose. *ubp3Δ* cells have lower amounts of Pho84 and Pho12 (Figure 3D). Further, upon shifting to low Pi for one hour, Pho84 and Pho12 increased in *ubp3Δ*. These data suggest that reduced Pho84 and Pho12 amounts in *ubp3Δ* are because of increased Pi, and not due to altered Pho regulon function itself (Figure 3D). These data clarify earlier observations from *ubp3Δ* which noted reduced Pho proteins (Isasa et al., 2015). Therefore, *ubp3Δ* constitutively have higher Pi, and likely a consequent decrease in Pho proteins.

We next asked if the increased Pi in *ubp3Δ* is because of altered G6P allocations towards different end-points, particularly trehalose synthesis, which can be a major node of Pi restoration (Gupta et al., 2019; Van Heerden et al., 2014). We assessed the contribution of increased trehalose synthesis towards the high Pi in *ubp3Δ*, by estimating Pi levels in the absence of trehalose 6-phosphate phosphatase (Tps2), which catalyzes the Pi-releasing step in trehalose synthesis. Notably, loss of Tps2 in *ubp3Δ* decreased Pi (Figure 3E). There was no additive difference in Pi between *tps2Δ* and *ubp3Δtps2Δ* (Figure 3E). As an added control, we assessed trehalose in WT and *ubp3Δ* in the absence of Tps2, and found no difference (Figure 3-figure supplement 1B). Therefore, increasing G6P flux towards trehalose biosynthesis is a major source of the increased Pi in *ubp3Δ*.

Since the major GAPDH isozymes, Tdh2 and Tdh3, are reduced in *ubp3Δ*, we directly asked if reducing GAPDH can decrease Pi consumption and increase Pi. To assess this, we generated *tdh2Δtdh3Δ* cells, which exhibit a growth defect, but are viable, permitting further analysis. *tdh2Δtdh3Δ* had higher Pi in high glucose (Figure 3F). Expectedly, we observed a decrease in ethanol in *tdh2Δtdh3Δ* (Figure 3-figure supplement 1C), along with an accumulation of F1,6 BP and G3P, and decreased 3PG and PEP (Figure 3-figure supplement 1D). However, G6P and trehalose levels between WT and *tdh2Δtdh3Δ* were comparable (Figure 3-figure supplement 1D, E). These data sugest that unlike in *ubp3Δ*, the increased Pi in *tdh2Δtdh3Δ* comes mainly from decreased Pi consumption (GAPDH step). To further assess the role of reduced glycolytic flux in increasing Pi (*ubp3Δ* and *tdh2Δtdh3Δ*), we measured the Pi in these cells growing in a gluconeogenic medium – 2%, ethanol. In this scenario, the GAPDH catalyzed reaction will be reversed, converting 1,3BPG to G3P, which should release and not consume Pi. Compared to WT, the Pi levels decreased in *ubp3Δ* and *tdh2Δtdh3Δ* (Figure 3G), suggesting that the changes in Pi in these mutants is driven by the relative change in Pi release vs consumption.

These results collectively indicate that the combined effect of increased Pi release coming from trehalose synthesis, and decreased Pi consumption from reduced GAPDH increase Pi levels in *ubp3Δ*.

### Mitochondrial Pi availability correlates with mitochondrial activity in *ubp3Δ*

We therefore wondered if this observed Pi increase from the combined rewiring of glucose metabolism resulted in more Pi becoming accessible to the mitochondria. This would effectively result in Pi budgeting between cytosolic glycolysis and mitochondria based on the relative flux in different arms of glucose metabolism, by increasing Pi availability for the mitochondria (Figure 4A). If this were indeed so, a prediction would be that the pool of Pi in the mitochondria of *ubp3Δ* cells would be higher than WT cells, while the opposite would be expected in the cytosolic fraction of these cells, due to increased Pi in the mitochondria. To test if this were so, we first measured the cytosolic pools of Pi in WT and *ubp3Δ.* The cytosolic fraction was isolated (extensive experimental details are in the SI appendix) from WT and *ubp3Δ* cells, and Pi levels in this fraction were estimated (Figure 4B, Figure 4-figure supplement 1B). We observed significantly reduced Pi in the cytosolic fraction in *ubp3Δ* cells. Since total cellular Pi amounts is higher in *ubp3Δ* cells (Figure 3), a decreased cytolosolic Pi would be consistent with greater transport of Pi from cytosol to other organelles such as vacuole (where it is stored as polyphosphate) and/or mitochondria. Therefore, we next asked if the mitochondria in *ubp3Δ* have correspondingly increased Pi. Mitochondria were isolated by immunoprecipitation from WT and *ubp3Δ*, the isolation efficiency was analyzed (Figure 4-figure supplement 1A), and relative Pi levels compared. Pi levels were normalised to Idh1 (isocitrate dehydrogenase) in isolated mitochondria, since Idh1 protein levels did not decrease in *ubp3Δ* (Figure 4-figure supplement 1B), and consistent with (Isasa et al., 2015). Mitochondrial Pi was higher in *ubp3Δ* (Figure 4B). We next asked if the increased mitochondrial activity in *ubp3Δ* is a consequence of high Pi. If this were so, *tdh2Δtdh3Δ* should partially phenocopy *ubp3Δ* with respect to mitochondrial activity. Consistently, *tdh2Δtdh3Δ* have higher mitotracker intensity as well as Cox2 levels compared to WT (Figure 4-figure supplement 1C, Figure 4C). Also consistent with this, a high basal OCR in *tdh2Δtdh3Δ* was observed (Figure 4D), together indicating high mitochondrial activity in *tdh2Δtdh3Δ*, which is comparable to *ubp3Δ*.

**Figure 4:**
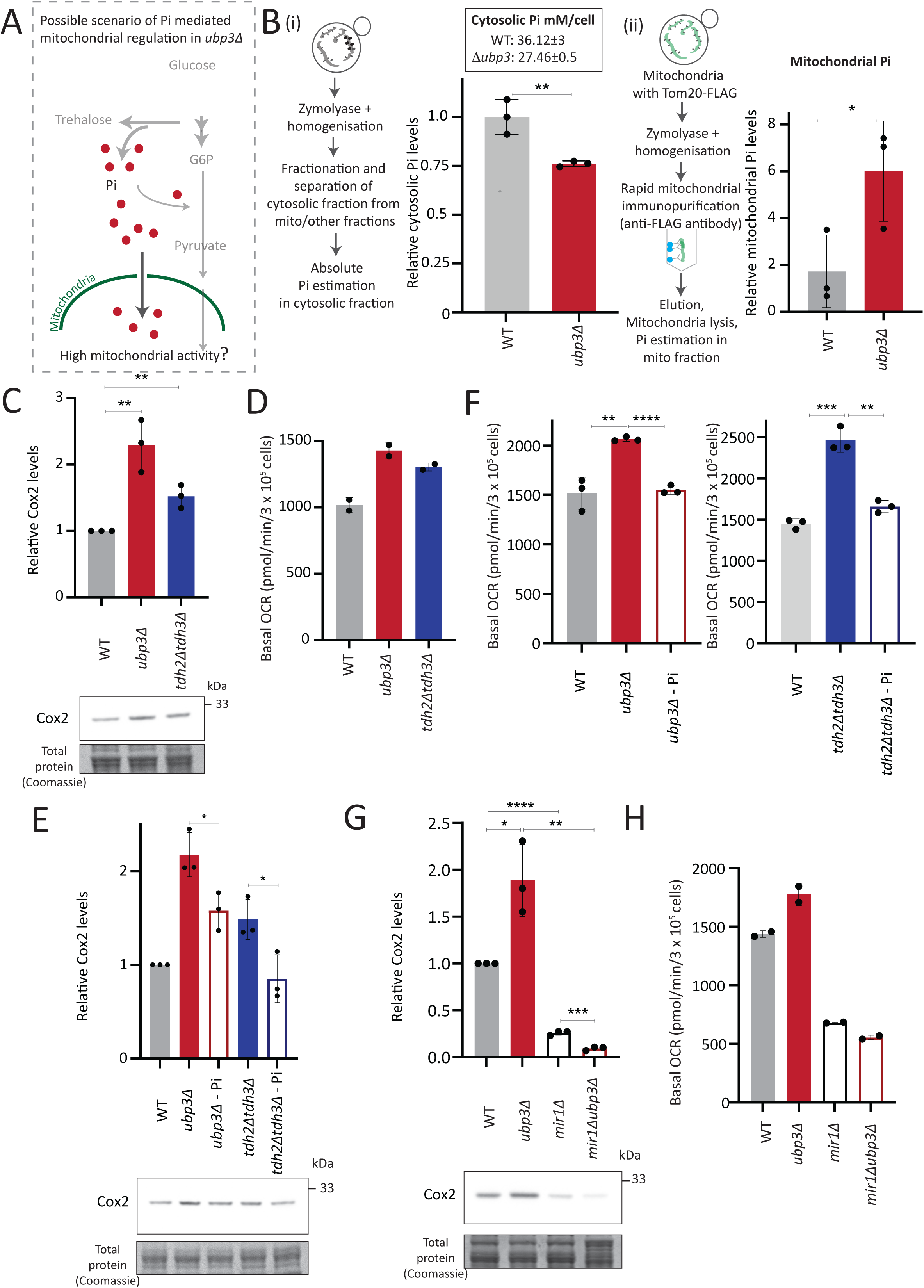
Mitochondrial Pi availability correlates with mitochondrial activity in *ubp3Δ*. A) A hypothetical mechanism of cytosolic free Pi controling mitochondrial activity by regulating mitochondrial Pi availability. B) Cytosolic and mitochondrial Pi amounts in WT vs *ubp3Δ*. The cytosolic fraction was isolated by centrifugation (see SI appendix), and in separate experiments, mitochondria were isolated by immunoprecipitation from WT and *ubp3Δ* and mitochondrial Pi estimated. (i) Cytosolic Pi levels (relative as well as absolute) and (ii) mitochondrial Pi levels (normalised to Idh1) are shown. Data represent mean ± SD from three biological replicates (n=3) respectively for the cytosolic and mitochondrial measurements. Also see Figure 4-figure supplement 1A, B. C) Cox2 protein in *tdh2Δtdh3Δ*. WT, *ubp3Δ,* and *tdh2Δtdh3Δ* were grown in high glucose and Cox2 protein was estimated. A representative blot (out of three biological replicates, n=3) and their quantifications are shown. Data represent mean ± SD. D) Basal OCR levels in *tdh2Δtdh3Δ*. WT, *ubp3Δ,* and *tdh2Δtdh3Δ* were grown in high glucose and basal OCR was measured from two independent experiments (n=2). Data represent mean ± SD. Also see Figure 4-figure supplement 1C. E) Comparative Pi amounts and Cox2 levels in *ubp3Δ*, *tdh2Δtdh3Δ,* WT cells. WT cells were grown in high glucose, *ubp3Δ* and *tdh2Δtdh3Δ* were grown in high glucose and low Pi, and Cox2 protein was estimated. A representative blot (out of three biological replicates, n=3) and their quantifications are shown. Data represent mean ± SD. Also see figure Figure 4-figure supplement 1D, F. F) Pi amounts and basal OCR in *ubp3Δ* and *tdh2Δtdh3Δ* vs WT cells. WT cells were grown in high glucose, *ubp3Δ* and *tdh2Δtdh3Δ* were grown in high glucose and low Pi, and basal OCR was measured from three independent experiments (n=3). Data represent mean ± SD. G) Effect of loss of mitochondrial Pi transporter Mir1 on Cox2protein. WT, *ubp3Δ*, *mir1Δ,* and *mir1Δubp3Δ* were grown in high glucose and Cox2 amounts compared. A representative blot (out of three biological replicates, n=3) and their quantifications are shown. Data represent mean ± SD. H) Relationship of mitochondrial Pi transport and basal OCR in WT vs *ubp3Δ*. WT, *ubp3Δ*, *mir1Δ,* and *mir1Δubp3Δ* cells were grown in high glucose and basal OCR was measured from two independent experiments (n=2). Data represent mean ± SD. Data information: *p<0.05, **p<0.01, ****p<0.0001.

We next asked if higher Pi is necessary to increase mitochondrial activity in *ubp3Δ* and *tdh2Δtdh3Δ*. Since glycolysis is defective in *ubp3Δ* and *tdh2Δtdh3Δ*, it is necessary to distinguish the effect of high Pi vs. only the effect of low glycolysis in activating mitochondria. Logically, if decreased glycolysis (independent of Pi) is sufficient to activate mitochondria, bringing down the Pi levels in *ubp3Δ* to that of WT should not affect mitochondrial activity. Notably, *ubp3Δ* grown in low (1 mM) Pi have Pi levels similar to WT in standard (normal Pi) medium (Figure 4-figure supplement 1D). *ubp3Δ* grown in low Pi also had decreased ethanol, suggesting reduced glycolysis (Figure 4-figure supplement 1E). Therefore, we used this condition to further understand the role of Pi in inducing mitochondrial activity. Mitotracker fluorescence decreased in both *ubp3Δ* and *tdh2Δtdh3Δ* in low Pi (Figure 4-figure supplement 1F). Note: basal mitotracker fluorescence in WT also decreases in low Pi, which is consistent with a required role of Pi for mitochondrial activity (Figure 4-figure supplement 1F). Similarly, Cox2 levels were reduced in both *ubp3Δ* and *tdh2Δtdh3Δ* (Figure 4E). Consistent with both reduced mitotracker intensity and Cox2 levels, basal OCR also decreased in both *ubp3Δ* and *tdh2Δtdh3Δ* in low Pi (Figure 4F). As an additional control, we used Rho0 strains (which have no mitochondrial DNA and therefore lack functional ETC) to compare basal OCR. The expectation in these strains is that basal OCR will not change if Pi changes. Consistently, we did not observe any significant difference in basal OCR in WT, *ubp3Δ* and *ubp3Δ* in low Pi in a Rho0 strain background (which lacks mitochondrial DNA) (Figure 4-figure supplement 1G). These data collectively suggest that high intracellular Pi is necessary to increase mitochondrial activity in *ubp3Δ* and *tdh2Δtdh3Δ*.

Next, we asked how mitochondrial Pi transport regulates mitochondrial activity. Mir1 and Pic2 are mitochondrial Pi transporters, with Mir1 being the major Pi transporter (Murakami et al., 1990; Zara et al., 1996). We first limited mitochondrial Pi availability in *ubp3Δ* by knocking-out *MIR1*. In *mir1Δ*, the increased Cox2 observed in *ubp3Δ* was no longer observed (Figure 4G). Consistent with this, we observed no further increase in the basal OCR in *mir1Δubp3Δ* compared to *mir1Δ* (Figure 4H). Furthermore, *mir1Δ* showed decreased Cox2 as well as basal OCR even in WT cells (Figure 4G, Figure 4H). These data together suggest that mitochondrial Pi transport is critical for increasing mitochondrial activity in *ubp3Δ*, and in maintaining basal mitochondrial activity even in high glucose.

As a control, no significant increase in Mir1 and Pic2 was observed in *ubp3Δ* (Figure 4-figure supplement 1H), suggesting that *ubp3Δ* do not increase mitochondrial Pi by merely increasing the Pi transporters, but rather by increasing available Pi pools.

Taken together, these data suggest the possibility that the altered Pi homeostais in *ubp3Δ* cells increases the mitochondrial Pi pool. This increased mitochondrial Pi pool correlates with increased mitochondrial activity. Decreasing mitochondrial Pi by either reducing total Pi, or by reducing mitochondrial Pi transport decreases mitochondrial activity.

### Mitochondrial Pi availability constrains mitochondrial activity under high glucose

So far, these data suggest that the cytosolic Pi available for the mitochondria can determine the extent of mitochondrial activity. Therefore, we further investigated if mitochondrial Pi allocation was a necessary constraint for glucose-mediated mitochondrial repression.

To test this, we first asked how important mitochondrial Pi transport was to switch to increased respiration. In WT yeast, low glucose or glycolytic inhibition will result in increased respiration (Broach, 2012). What happens therefore if we restrict mitochondrial Pi in this context? For this, we measured the basal OCR in WT and *mir1Δ* after switching from high (2%) to low (0.1%) glucose. We observed a significant increase in the basal OCR in WT but not in *mir1Δ* (Figure 5A). The alternate scenario is after glycolytic inhibition. We assessed the role of mitochondrial Pi in this context, by inhibiting glycolysis using 2-deoxyglucose (2DG). WT, but not *mir1Δ*, increased their OCR (respiration) upon a 1-hour treatment with 2DG (Figure 5B). Consistent with this, mitotracker fluorescence increased with an increase in 2DG in WT, but not in *mir1Δ* (Figure 5-figure supplement 1A). We further asked if the mitochondrial Pi transporter itself glucose repressed, and therefore assessed Mir1 amounts in high and low glucose. Mir1 levels are higher upon a shift to low glucose, and in cells grown in 2% ethanol, suggesting that Mir1 is glucose repressed (Figure 5C, Figure 5-figure supplement 1B). These data suggest that mitochondrial Pi transport is necessary for increasing mitochondrial activity after glucose derepression.

**Figure 5:**
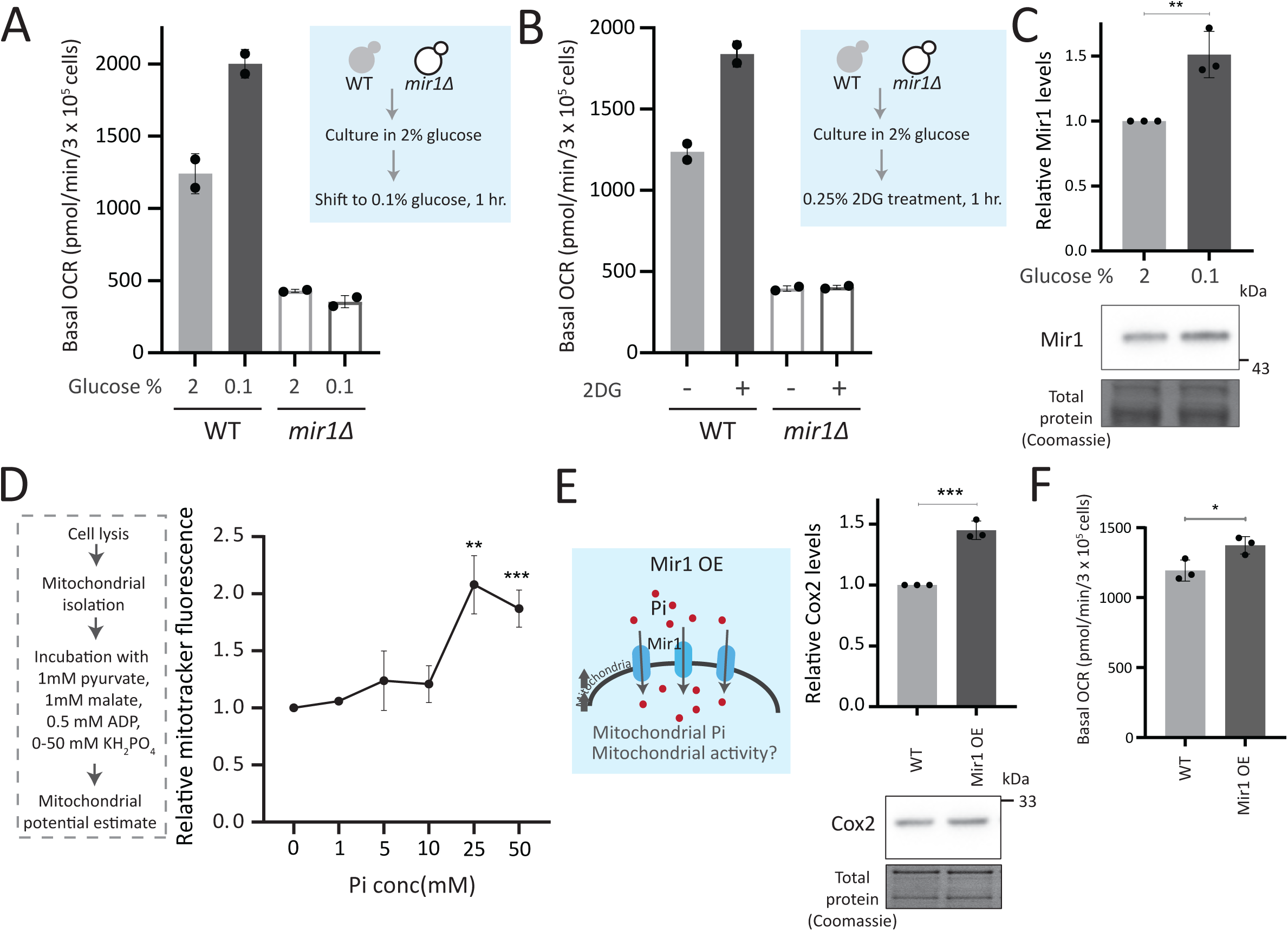
Mitochondrial Pi availability constrains mitochondrial activity under high glucose. A) Relationship of mitochondrial Pi transport and respiration after glucose removal. WT and *mir1Δ* cells were cultured in high (2%) glucose and shifted to low (0.1%) glucose for 1 hour. The normalized basal OCR, from two independent experiments (n=2) are shown. Data represent mean ± SD. B) Requirement of mitochondrial Pi transport for switch to respiration upon glycolytic inhibition by 2DG. WT and *mir1Δ* cells were cultured in high glucose and treated with or without 0.25% 2DG for 1 hour. Basal OCR was measured from two independent experiments (n=2). Data represent mean ± SD. Also see Figure 5-figure supplement 1A. C) Glucose-dependent regulation of Mir1. Cells (with Mir1-HA) were grown in high glucose and shifted to low glucose (0.1% glucose) for 1 hour, and Mir1 levels compared. A representative blot (out of three biological replicates, n=3) and their quantifications are shown. Data represent mean ± SD. Also see Figure 5-figure supplement 1B. D) Increasing Pi concentrations and mitochondrial activity in isolated mitochondria. Mitochondria were isolated from WT cells grown in high glucose, incubated with 1 mM pyruvate, 1 mM malate, 0.5 mM ADP and 0-50 mM KH_2_PO_4_. The mitochondrial activity was estimated by mitotracker fluorescence intensity, and intensities relative to the sample with 0 mM KH_2_PO_4_ is shown. Data represent mean ± SD from three biological replicates (n=3). E) Effect of overexpressing Mir1 on Cox2 protein. WT (containing empty vector) and Mir1 overexpressing (Mir1OE) cells were grown in high glucose and Cox2 levels were estimated. A representative blot (out of three biological replicates, n=3) and their quantifications are shown. Data represent mean ± SD. Also see figure S5F. F) Effect of overexpressing Mir1 on basal OCR. The basal OCR in WT (containing empty vector) and Mir1OE in high glucose was measured from three independent experiments (n=3). Data represent mean ± SD. Data information: *p<0.05, **p<0.01, ***p<0.001.

We next asked whether just adding external Pi was sufficient to increase mitochondrial activity, when cells are in high glucose. In medium supplemented with excess Pi, the internal Pi increases as seen earlier (Figure 3C). Therefore, a simplistic assumption would be that the addition of external Pi to cells in high glucose would also increase mitochondrial Pi. However, an alternate possibility presents itself whererin since glycolytic flux is already high in glucose, supplementing Pi will continue to fuel glycolysis. Indeed, this was originally obseved by Harden and Young in 1908, where adding Pi increased fermentation (Harden and Young, 1906). In such a scenario, there could be an increase in the cytosolic Pi but not the mitochondrial Pi. We estimated the cytosolic and mitochondrial Pi in this condition where excess Pi was externally supplemented. Notably, cells grown in high Pi had increased cytosolic Pi, but decreased mitochondrial Pi (Figure 5-figure supplement 1C), without any changes in total mitochondria volume or amounts (Figure 5-figure supplement 1D). Furthermore, directly adding Pi to cells growing in high glucose also decreased basal OCR (Figure 5-figure supplement 1E), consistent with decreased mitochondrial Pi. These data indicate that in high glucose, simply supplementing Pi will not increase Pi access to the mitochondria, and instead results in an accumulation of Pi in the cytosol (Figure 5-figure supplement 1C). We therefore now asked, if we inhibit glycolytic flux and then supplement Pi, what would happen to mitochondrial activity. For this, we treated cells with 2DG, and subsequently added Pi and measured the OCR (Figure 5-figure supplement 1F). In this case, supplementing Pi increased the basal OCR (Figure 5-figure supplement 1F). Collectively, these data suggest that a combination of decreasing glycolysis and increasing Pi can together increase respiration. Next, in order to directly test mitochondrial activation based on external Pi availability, we isolated mitochondria, and estimated activity *in vitro* upon adding increasing Pi. Mitochondrial activity increased with increased Pi, with maximum activity observed with 25 mM Pi supplemented (Figure 5D). In a complementary experiment, we overexpressed the Mir1 transporter in wild-type cells, to increase Pi within mitochondria (Figure 5-figure supplement 1G). Mir1-OE cells have higher Cox2 levels and basal OCR (Figure 5E, 5F). Therefore, increasing Pi transport to mitochondria is sufficient to increase mitochondrial activity in high glucose.

Mitochondrial pyruvate transport is also required for mitochondrial respiration (Timoń-Gómez et al., 2013). We asked where Pi availability stands in a hierarchy of constraints for mitochondrial derepression, as compared to mitochondrial pyruvate transport. We measured the amounts of the Mpc3 subunit of the mitochondrial pyruvate carrier (MPC) complex (Bender et al., 2015; Timoń-Gómez et al., 2013). Mpc3 protein increases in *ubp3Δ* (Figure 5-figure supplement 1H). This also correlated with the unimpaired TCA cycle flux (Figure 2-figure supplement 2D) and the increased mitochondrial activity. Interestingly, in *ubp3Δ* grown in low Pi, Mpc3 further increased (Figure 5-figure supplement 1H), but as shown earlier this condition cannot increase OCR or mitochondrial activity (Figure 4F, Figure 4-figure supplement 1F). Basal Mpc3 levels decrease in *mir1Δ*, but upon shifting to 0.1% glucose, Mpc3 increases in both WT and *mir1Δ*, with higher levels in *mir1Δ* (Figure 5-figure supplement 1I). Therefore, even where Mpc3 is high (*ubp3Δ* in low Pi, and *mir1Δ* in low glucose), mitochondrial activity remains low if Pi is restricted (Figure 4F, 4H). There was also no decrease in basal OCR in *mpc3Δ* in high glucose, and the basal OCR increased to the same level as of WT after shifting to 0.1% glucose (Figure 5-figure supplement 1J). Since Mpc3 changes with mitochondrial Pi availability (Figure S5G, S5H), we also measured Mpc3 in the Mir1OE. No further changes in Mpc3 were observed in Mir1OE (Figure 5-figure supplement 1K), indicating that increasing mitochondrial Pi alone need not increase Mpc3. Overall, although Mpc3 levels corelate with decreased glycolysis (*ubp3Δ* - Figure 5-figure supplement 1H, *ubp3Δ* in low Pi-Figure 5-figure supplement 1H, low glucose - Figure 5-figure supplement 1I), increased Mpc3 alone cannot increase mitochondrial activity and respiration in the absence of adequate mitochondrial Pi.

Collectively, mitochondrial Pi availability constrains glucose-mediated mitochondrial repression. Increasing available pools of Pi to enter the mitochondria is sufficient to induce mitochondrial activity.

### Repression of mitochondrial respiration via Pi budgeting is conserved in Ubp3 mutants across diverse yeast genetic backgrounds

So far, we have identified a role for intracellular Pi budgeting as a constraint for mitochondrial activity under high glucose. These were all carried out using a robust, prototrophic yeast strain from a CEN.PK background. *S. cerevisiae* however, while Crabtree positive, have tremendous genetic diversity (Peter et al., 2018). We therefore asked if this mitochondrial repression through Pi budgeting (mediated by Ubp3 function) is conserved across other strains of *S. cerevsiae* as well. To test this, we generated Ubp3 deletion mutants in different genetic backgrounds of *S.* cerevsiae, including BY4742, W303 and Σ1278. In all these strain backgrounds, we observed a significant increase in mitotracker fluorescne intensity and Cox2 protein levels in *ubp3Δ* (Figure 6A, B). This suggests the role of Ubp3 as a regulator of mitochondrial repression, independent of the genetic background of the yeast (*S. cerevsiae*) strain. To further assess if loss of Ubp3 shows a concurrent increase in Pi levels, we measured the total Pi levels in WT and *ubp3Δ* in a W303 strain background. We observed a significant increase in total Pi levels in *ubp3Δ* cells in this strain (Figure 6C), similar to what we observed in the CEN.PK strain (Figure 3C). Finally, to test if the altered Pi budgeting regulates the mitochondrial activity in these cells, we measured the basal OCR in WT, *ubp3Δ* and *ubp3Δ* in low (1 mM) Pi medium, in the W303 strain background. Consistent with the increase in mitotracker fluorescence intensity and Cox2 protein levels, we observed a significant increase in the basal OCR in *ubp3Δ* cells (Figure 6D). This increase was not observed in *ubp3Δ* in a low Pi medium (Figure 6D). This suggests that the role of altered Pi budgeting in regulating mitochondrial respiration is conserved in other genetic backgrounds of *S. cerevisiae*.

**Figure 6:**
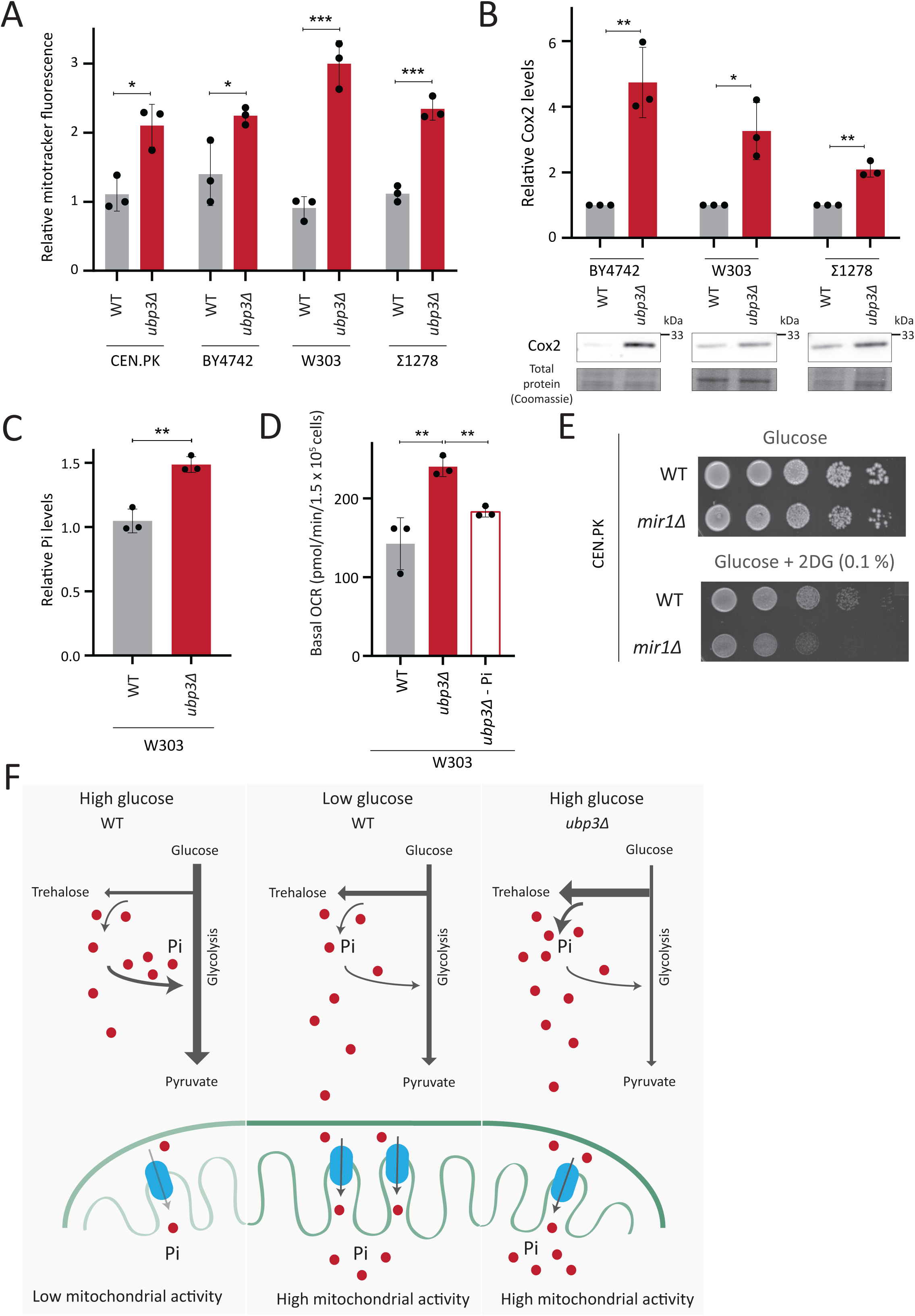
Repression of mitochondrial respiration via Pi budgeting is conserved in Ubp3 mutants across diverse yeast genetic backgrounds. A) Effect of loss of Ubp3 on mitochondrial membrane potential in different yeast strains. WT and *ubp3Δ* cells (in CEN.PK, BY4742, W303 and Σ1278 strains of *S. cerevisiae*) were grown in high glucose and relative mitochondrial membrane potential was measured. Data represent mean ± SD from three biological replicates (n=3). B) Effect of loss of Ubp3 on ETC complex IV subunit Cox2. WT and *ubp3Δ* (in BY4742, W303 and Σ1278 strains of *S. cerevisiae*) were grown in high glucose, and Cox2 was measured. A representative blot (out of 3 biological replicates, n=3) and their quantifications are shown. Data represent mean ± SD. C) Intracellular Pi levels in WT and *ubp3Δ* in W303 strain background. WT and *ubp3Δ* (in W303 strain background) were grown in high glucose and the total free phosphate (Pi) levels were estimated. Data represent mean ± SD from three biological replicates (n=3). D) Effect of low Pi on the basal OCR in WT and *ubp3Δ* cells in W303 strain background. WT cells were grown in high glucose and *ubp3Δ* were grown in high glucose and low Pi, and basal OCR was measured. Data represent mean ± SD (n=3). E) Requirement of mitochondrial Pi transport for growth after 2DG treatment. Shown are serial dilution growth assays in high glucose in the presence and absence of 0.1% 2DG, using WT and *mir1Δ* cells. The results after 40hrs incubation/30°C are shown. F) A model illustrating how mitochondrial Pi availability controls mitochondrial activity. In high glucose, the decreased Pi due to high Pi consumption in glycolysis, along with the glucose-mediated repression of mitochondrial Pi transporters, decreases mitochondrial Pi availability. This reduces mitochondrial activity. In low glucose, increased mitochondrial Pi transporters and lower glycolytic flux increases mitochondrial Pi, leading to enhanced mitochondrial activity. In *ubp3Δ* cells in high glucose, high trehalose synthesis and lower glycolytic flux results in an increase in Pi. This increases mitochondrial Pi availability and thereby the mitochondrial activity. Data information: *p<0.05, **p<0.01, ***p<0.001.

Finally, we asked how important mitochondrial Pi transport was for growth, under glycolytic inhibition. Consistent with the requirement for mitochondrial Pi transport to increase mitochondrial respiration upon glycolytic inhibition (Figure 5B), WT cells only exhibit a slightly decreased growth in the presence of 2DG (Figure 6E). In contrast, *mir1Δ* show a severe growth defect upon 2DG treatment (Figure 6E), revealing a synergetic effect of combining 2DG with inhibiting mitochondrial Pi transport. Therefore, the combined inhibition of glycolysis and mitochondrial Pi transport restricts the growth of glycolytic cells.

Collectively, our data suggests a conserved role for intracellular Pi bugeting in regulating mitochondrial repression in high glucose and the role of mitochondrial Pi transport in regulating adaptation for growth under glycolytic inhibition.

## Discussion

In this study, we highlight a role for Pi budgeting between cytosolic glycolysis and mitochondrial processes (which compete for Pi) in constraining mitochondrial repression (Figure 6F). Ubp3 controls this process (Figure 1), by maintaining the amounts of the glycolytic enzymes Pfk1 and GAPDH (Tdh2 and Tdh3) and thereby allows high glycolytic flux. At high fermentation rates, glycolytic enzymes levels at maximal activity maintain high glycolytic rates (effectively following zero-order kinetics), and therefore, changes in the enzyme levels will have a direct, proportionate effect on flux (Grigaitis and Teusink, 2022). The loss of Ubp3 decreases glycolytic flux, resulting in a systems-level, mass-action based rewiring of glucose metabolism where more G6P is routed towards trehalose synthesis and PPP (Figure 2, Figure 6F). Indeed, inhibiting just phosphofructokinase can reroute glucose flux from glycolysis to PPP (Hollinshead et al., 2016; Miyazawa et al., 2017; Yi et al., 2012), and our study now permits contextualized interpretations of these results. Such reallocations of glucose flux will collectively increase overall Pi, coming from the combined effect of increased Pi release from trehalose synthesis, and decreased Pi consumption via reduced GAPDH. This altered intracellular Pi economy increases Pi pools available to mitochondria, and increasing mitochondrial Pi is necessary and sufficient to increase respiration in high glucose (Figure 4, Figure 5, Figure 6). Mitochondrial Pi transport maintains mitochondrial activity in high glucose, and increases mitochondrial activity in low glucose (Figure 4, Figure 5). Finally, the mitochondrial Pi transporter Mir1 itself decreases in high glucose (Figure 5). Therefore, this glucose-dependent repression of Mir1 also restricts mitochondrial Pi availability (and thereby activity) in high glucose.

Traditionally, loss-of-function mutants of metabolic enzymes are used to understand metabolic state regulation. This approach negates nuanced investigations, since metabolic enzymes are often essential for viability. Furthermore, metabolic pathways have multiple, contextually regulated nodes, through which cells maintain their metabolic state. Therefore, alternate approaches to identify global regulators of metabolic states (as opposed to single enzymes), might uncover ways via which multiple nodes are simultaneously tuned, and can reveal unanticipated systems-level principles of metabolic state rewiring. In this study of glucose-mediated mitochondrial repression, the loss of the DUB Ubp3 decreases glycolytic flux by reducing the enzymes at two critical nodes in the pathway - Pfk1 and GAPDH. Unlike loss of function mutants, a reduction in amounts will only rewire metabolic flux. By ‘hitting’ multiple steps in glycolysis simultaneously, *ubp3Δ* have decreased Pi consumption, as well as increased Pi release. Such a cumulative phenomenon reveals more than inhibiting only GAPDH, where increased Pi comes only from reduced Pi consumption, and not from increased trehalose biosynthesis. Our serendipitous identification of a regulator which regulates multiple steps in glucose metabolism to change the metabolic environment, now suggests a general basis of mitochondrial regulation that would have otherwise remained hidden. Separately, finding the substrates of Ubp3 and whether Ubp3 directly regulates glycolytic enzymes are exciting future research questions requiring concurrent innovations in accessible chemical-biological approaches to study DUBs.

Because phosphates are ubiquitous, it is challenging to identify hierarchies of Pi-dependent processes in metabolic state regulation (Gupta and Laxman, 2021). Phosphate transfer reactions are the foundation of metabolism, driving multiple, thermodynamically unfavorable reactions (Kamerlin et al., 2013; Westheimer, 1987). Contextually, the laws of mass action predict that the relative rates of these reactions will regulate overall Pi balance, and contrarily the Pi allocation to Pi-dependent reactions will determine reaction rates (Gupta and Laxman, 2021; Van Heerden et al., 2014). Additionally, cells might control Pi allocations for different reactions via compartmentalizing Pi in organelles, to spatially restrict Pi availability (Booth and Guidotti, 1997; Solesio et al., 2021; Vila et al., 2022). Our data collectively suggest a paradigm where the combination of factors regulates mitochondrial Pi and thereby activity. In glycolytic yeast cells growing in high glucose, mitochondrial Pi availability becomes restricted due to higher utilization of Pi in glycolysis compared to mitochondria. Consistent with this, a rapid decrease in Pi upon glucose addition has been observed (Hohmann et al., 1996; Koobs, 1972; Rodríguez-Enríquez et al., 2001). Interesting, *in vitro* studies with isolated mitochondria from tumor cells also find that decreasing Pi levels decreases respiration (Rodríguez-Enríquez et al., 2001), which would be consistent with this scenario. Further, supplementing Pi correlates with decreased mitochondrial repression in tumors (Brin and McKee, 1956; Koobs, 1972). By increasing Pi through a systems-level rewiring of glucose metabolism (such as in *ubp3Δ* cells), cells can collectively increase mitochondrial access to Pi. This Pi budgeting determines mitochondrial activity. Supplementing Pi under conditions of low glycolysis (where mitochondrial Pi transport is enhanced), as well as directly supplementing Pi to isolated mitochondria, increases respiration (Figure 5, Figure 5-figure supplement 1). Notably, this increased respiration does not happen upon directly supplementing Pi to highly glycolytic WT cells, where the Pi increases in cytosol, without increasing mitochondrial Pi (Figure 5-figure supplement 1C). Therefore, in order to derepress mitochondria, a combination of increased Pi along with decreased glycolysis is required. An additional systems-level phenomenon that might regulate Pi transport to the mitochondria is the decrease in cytosolic pH upon decreased glycolysis (Dechant et al., 2010; Orij et al., 2011). The cytosolic pH in highly glycolytic cells is ∼7, and decreasing glycolysis results in cytosolic acidification (Dechant et al., 2010; Orij et al., 2011). Therefore, under conditions of decreased glycolysis (2DG treatment, deletion of Ubp3, and decreased GAPDH activity), cytosolic pH becomes acidic. Since mitochondrial Pi transport itself is dependent on the proton gradient, a low cytosolic pH would favour mitochondrial Pi transport (Hamel et al., 2004). Therefore, under conditions of decreased glycolysis (2DG treatment, or loss of Ubp3, or decreased GAPDH activity), where cytosolic pH would be acidic, increasing cytosolic Pi might indirectly increase mitochondria Pi transport, thereby leading to increased respiration. Alternately, increasing mitochondrial Pi transporter amounts can achieve the same result, as seen by overexpressing Mir1 (Figure 5). A similar observation has been reported in *Arabidopsis*, reiterating an evolutionarily conserved role for mitochondrial Pi in controlling respiration (Jia et al., 2015). Relatedly, glycolytic inhibition can suppress cell proliferation in Warburg-positive tumors (O’Neill et al., 2019; Pelicano et al., 2006), or inflammatory responses (Soto-Heredero et al., 2020). However, these cells survive by switching to mitochondrial respiration (Lu et al., 2015; Shiratori et al., 2019), requiring alternate approaches to prevent their proliferation (Cheng et al., 2012). Inhibiting mitochondrial Pi transport in combination with glycolytic inhibition could restrict the proliferation of Warburg/Crabtree positive cells. It is important to highlight that our experiments, whether involving Pi supplementation or Pi limitations, maintain the cellular Pi concentration within the millimolar range, and are conducted within a short timeframe (∼ 1 hour). This differs significantly from Pi starvation studies, where cells are subjected to prolonged and complete Pi deprivation. In those contexts, cells trigger extensive metabolic adaptations in order to sustain available Pi pools including an increase in mitochondrial membrane potential which can be independent of respiration (Ouyang et al., 2024).

Since its discovery in the 1920s, the phenomenon of accelerated glycolysis with concurrent mitochondrial repression has been intensely researched. Yet, the biochemical constraints for glucose-mediated mitochondrial repression remains unresolved. One hypothesis suggests that the availability of glycolytic intermediates might determine the extent of mitochondrial repression. F1,6BP inhibits complex III and IV of the electron transport chain (ETC) in Crabtree positive yeast (Diaz-Ruiz et al., 2011; Díaz-Ruiz et al., 2008; Hammad et al., 2016; Lemus et al., 2018). Similarly, the ratio between G6P and F1,6BP regulates the extent of mitochondrial repression (Díaz-Ruiz et al., 2008; Lemus et al., 2018). Although G6P/F6P accumulates in *ubp3Δ* (Figure 2C), this is not the case in *tdh2Δtdh3Δ* (GAPDH mutant) (Figure 3-figure supplement 1D), suggesting that G6P/F6P accumulation in itself is not the criterion to increase mitochondrial activity. Separately, the competition for common metabolites/co-factors between glycolysis and respiration (such as ADP, Pi or pyruvate) could drive this phenomenon (Diaz-Ruiz et al., 2011; Koobs, 1972). Here we observe that Mpc3 mediated mitochondrial pyruvate transport alone cannot increase respiration. An additional consideration is the possible contribution of changes in ADP in regulating mitochondrial activity, where the use of ADP in glycolysis might limit mitochondrial ADP. Therefore, when Pi changes as a consequence of glycolysis, it coud be imagined that a change in ADP balance can coincidentally occur. However, prior studies show that even though cytosolic ADP decreases in the presence of glucose, this does not limit mitochondrial ADP uptake, or decrease respiration, due to the very high affinity of the mitochondrial ADP transporter (Diaz-Ruiz et al., 2011; Rodríguez-Enríquez et al., 2001). These collectively reiterate the importance of Pi access and transport to mitochondria in constraining mitochondrial respiration. Indeed, this interpretation can also contextually explain observations from other model systems where mitochondrial Pi transport seems to regulate respiration (Scheibye-Knudsen and Quistorff, 2009; Seifert et al., 2015).

We parsimoniously suggest that Pi access to the mitochondria as a key constraint for mitochondrial repression under high glucose. In a hypothetical scenario, a single-step event in evolution, reducing mitochondrial Pi transporter amounts, and/or increasing glycolytic flux (to deplete cytosolic Pi), will result in whole-scale metabolic rewiring to repress mitochondria. More elaborate regulatory events can easily be imagined as subsequent adaptations to enforce mitochondrial repression. Given the central role played by Pi, something as fundamental as access to Pi will constrain mitochondrial repression. Concurrently, the rapid incorporation of Pi into faster glycolysis can give cells a competitive advantage, while also sequestering Pi in the form of usable ATP. Over the course of evolution, this could conceivably drive other regulatory mechanisms to enforce mitochondrial repression, leading to the currently observed complex regulatory networks and signaling programs observed in the Crabtree effect, and other examples of glucose-dependent mitochondrial repression.

## Materials and methods

### Statistics and graphing

Unless otherwise indicated, statistical significance for all indicated experiments were calculated using unpaired Student’s t-tests (GraphPad prism 9.0.1). Graphs were plotted using GraphPad prism 9.0.1

### Yeast strains, media, and growth conditions

A prototrophic CEN.PK strain of *Saccharomyces cerevisiae* (WT) (J.P. van Dijken et al., 2000) was used unless mentioned otherwise. Strains are listed in appendix table S1. Gene deletions, chromosomal C-terminal tagged strains were generated by PCR mediated gene deletion/tagging (Longtine et al., 1998). Mitochondria targeted Mneon strain (Mito-mNeon green) is described in (Dua et al., 2022). The cox2-62 strain is described in (Bonnefoy et al., 2001). Media compositions, growth conditions and CRISPR-Cas9 based mutagenesis are described in extended methods (SI appendix).

### Mitotracker fluorescence

Mitotracker fluorescence was measured using Thermo Varioscan^TM^ LUX multimode plate reader (579/599 excitation/emission). Detailed protocol is described in extended methods. Mitotracker fluorescence were normalized using OD600 of each sample and relative fluorescence intensity calculated.

### Protein extraction and Western blotting

Total protein was precipitated, extracted using trichloroacetic acid (TCA) as described earlier (Vengayil et al., 2019). Blots were quantified using ImageJ software. Detailed protocol is described in extended methods.

### Basal OCR measurement

The basal OCR was measured using Agilent Seahorse XFe24 analyzer. Basal OCR readings were normalized for cell number (using OD_600_ of samples) in each well. The detailed methods are described in extended methods.

### Mitochondrial volume estimation

High-resolution 3D fluorescence experiments were performed on an inverted confocal laser scanning microscope (Carl Zeiss LSM 780 or Olympus FV3000). For each imaging field of view, sequential z-stacks were acquired for each excitation channel. 488 nm laser excitation for mNeonGreen and 561 nm laser excitation for Mitotracker CMXros dye were used respectively. Images taken were deconvolved and analyzed further in ImageJ software with custom-written routines. Mitochondria segmentation and quantification was done using the Mitochondria Analyzer plugin (Chaudhry et al., 2019) in Imagej. For visualization, maximum intensity projection of 3D images was used.

### RNA extraction and RT-qPCR

The RNA extraction was done using the hot phenol extraction method as described in (Vengayil et al., 2019). The isolated RNA was DNase treated, and used for cDNA synthesis. Superscript III reverse transcriptase enzyme (Invitrogen) was used for cDNA synthesis and RT-qPCR was performed using KAPA SYBR FAST qRT PCR kit (KK4602, KAPA Biosystems). Taf10 was used as a control for normalisation and the fold change in mRNA levels were calculated by 2^-ddct method.

### ATP, ethanol and Pi measurements

ATP levels were measured by ATP estimation kit (Thermo Fisher A22066). Ethanol concentration in the medium was estimated using potassium dichromate based assay described in (Sriariyanun et al., 2019) with modifications. Pi was estimated using a malachite green phosphate assay kit (Cayman chemicals, 10009325). Detailed sample collection and assay protocols are described in extended methods.

### Metabolite extraction and analysis by LC-MS/MS

The steady state levels and relative ^13^C label incorporation into metabolites were estimated by quantitative LC-MS/MS methods as described in (Walvekar et al., 2018). Detailed methodology is extensively described in extended methods. Peak area measurements are listed in supplement file 1.

### Mitochondrial isolation

Mitochondria was isolated by immunoprecipitation as described in (Chen et al., 2017; Liao et al., 2018) with modifications. The detailed protocol is described in extended methods (SI appendix). The eluted mitochondria were used in malachite green assay for Pi estimation, boiled with SDS-glycerol buffer for western blots or incubated with mitotracker CMXROS with mitochondria-activation buffer for mitotracker assays.

### Cytosolic fraction isolation

Cytosolic fraction was isolated from spheroplasts by centrifugation as described in detail in S1 appendix. The total protein amounts in the cytosolic fraction was estimated by BCA protein estimation assay and the Pi levels were estimated by malachite green assay.

## Data availability

The LC-MS/MS peak area for all the metabolites can be found in supplement file 1. This study includes no data deposited in external repositories.

## Acknowledgements

We thank all SL lab members, Benjamin Tu, Anand Bachhawat and Vijay Jayaraman for comments and suggestions. We acknowledge the NCBS/inStem/CCAMP mass spectrometry facility for LC-MS/MS support. We thank Aayushee Khanna for help with experiments and Gaurav Singh for help with microscopy. V.V acknowledges funding support from DST-INSPIRE fellowship (IF170236) from the Department of Science and Technology (DST), Govt. of India, S.N and SA acknowledges intramural funding from the DBT-inStem PhD program. S.L acknowledges support from the DST SERB CRG grant CRG/2019/004772, a DBT-Wellcome India Alliance Senior Fellowship (IA/S/21/2/505922), and institutional support from inStem.

## Supplement figure legends

**Figure 1-figure supplement 1:**
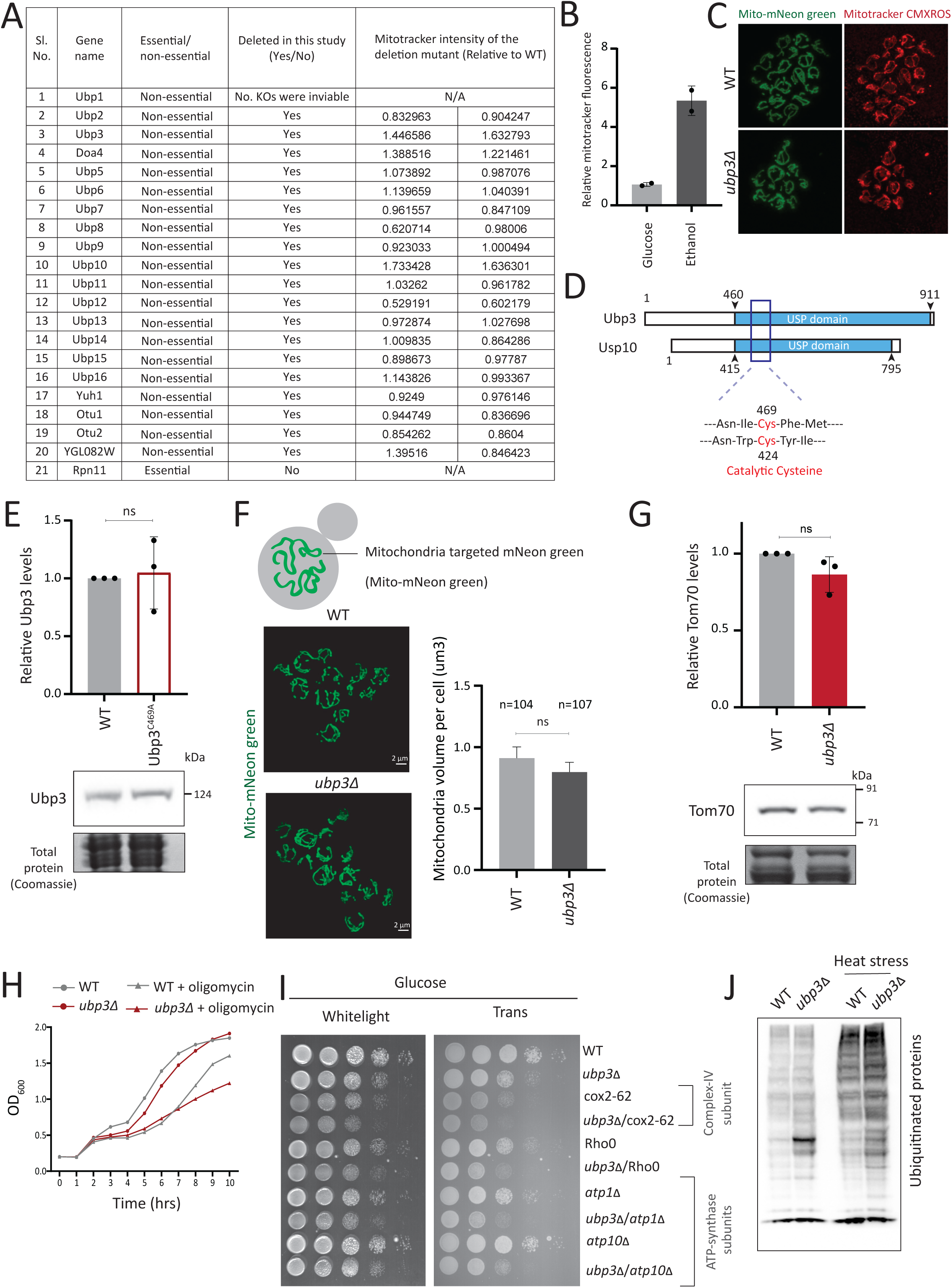
DUB screen details and further characterization of Ubp3 functions. A) A list of the known/identified *S. cerevisiae* deubiquitinase enzymes is shown. The DUBs which were used in this study, and the mitotracker intensity of two biological replicates (n=2) relative to WT are shown. B) WT cells grown in a respiratory, ethanol medium and mitotracker fluorescence intensity compared to glucose medium. WT cells were grown inhigh glucose medium (2% glucose) and ethanol medium (2% ethanol), and the mitochondrial membrane potential was measured. Data from two independent experiments (n=2) is shown. Data represent mean ± SD. C) Representative images of WT and *ubp3Δ* cells treated with mitotracker red CMXROS. D) Schematic representation of the conserved domains of Ubp3 and its mammalian ortholog Usp10. The catalytic cysteine of Usp10 and Ubp3 is highlighted. E) The catalytically inactive mutant of Ubp3, Ubp3^C469A^ does not have altered Ubp3 protein levels. WT and Ubp3^C469A^ cells containing endogenous Ubp3 tagged with 3xFLAG (C terminus) were grown in high glucose and the protein levels of Ubp3 were measured. A representative blot (out of three biological replicates, n=3) and their quantification are shown. Data represent mean ± SD. F) The total mitochondrial volume does not change in *ubp3Δ* cells. WT and *ubp3Δ* cells with mitochondria targeted mNeon green were imaged, and the total mitochondrial volume per cell (n=104 for WT, n=107 for *ubp3Δ*) was calculated. A representative image and quantification are shown. Data represent mean ± SD. G) Tom70 protein amounts do not change in *ubp3Δ* cells. WT and *ubp3Δ* cells containing endogenous Tom70 tagged at the C terminus with a 3xFLAG epitope were grown in high glucose and the protein levels of Tom70 were measured by western blot. A representative blot (out of three biological replicates, n=3) and their quantification are shown. Data represent mean ± SD. H) Dependence of *ubp3Δ* on mitochondrial respiration. WT and *ubp3Δ* cells were grown in high glucose with or without mitochondrial inhibitor oligomycin (25 µM), and the OD_600_ was measured at different time points. Data represent mean ± SD (n=2). I) Dependence of *ubp3Δ* on functional mitochondrial respiration. Ubp3 deletions were done in respiration defective strains -Rho0, cox2-62, *atp1Δ* and *atp10Δ*, and serial dilution growth assay was done in high glucose. The results after 24 hours of incubation at 30°C are shown. J) Changes in global ubiquitination state in WT and *ubp3Δ.* WT and *ubp3Δ* cells were grown in high glucose with or without heat stress (42°C/1 hour), and the changes in global total ubiquitination was estimated by western blotting using an anti-ubiquitin antibody.

**Figure 2-figure supplement 1:**
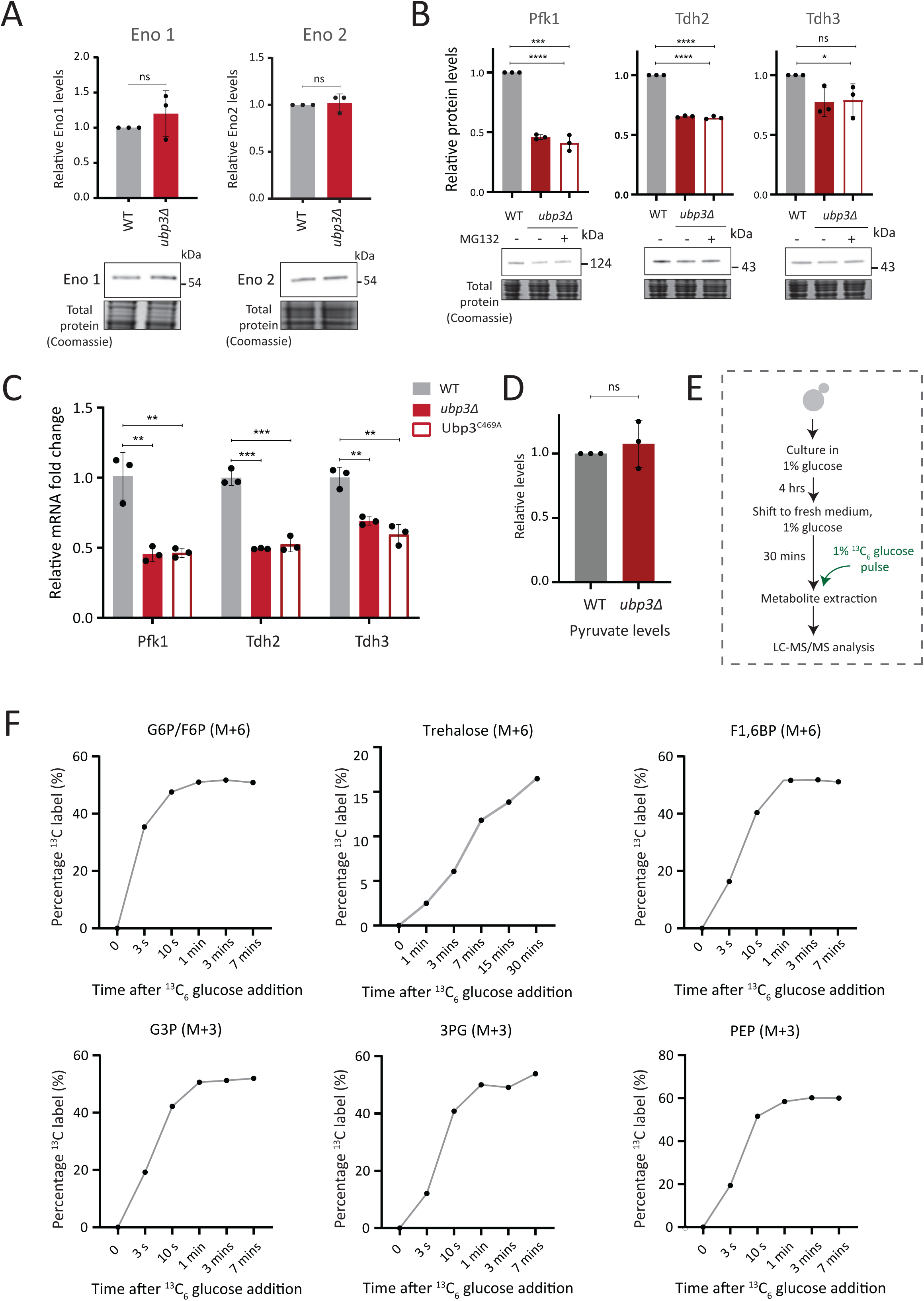
Further estimates of glycolytic enzymes and flux. A) Loss of Ubp3 does not change the protein levels of enolase isozymes Eno1 and Eno2. WT and *ubp3Δ* cells containing Eno1 and Eno2 C terminus endogenously tagged with a 3x FLAG epitope, were grown in high glucose and protein amounts were estimated. A representative blot (out of three biological replicates, n=3) is shown in the lower panel, quantifications shown in the upper panel. Data represent mean ± SD. B) The decreased levels of glycolytic enzymes in *ubp3Δ* cells is not because of proteasomal degradation. WT and *ubp3Δ* were grown in high glucose and *ubp3Δ* cells were treated with or without MG132 (100 μM) for 30 minutes. Pfk1, Tdh2, and Tdh3 levels were measured by western blot using an anti-FLAG antibody. A representative blot (out of three biological replicates, n=3) and their quantification are shown. Data represent mean ± SD. C) Loss of Ubp3 results in decreased transcription of PFK1, TDH2 and TDH3 genes. WT, *ubp3Δ* and Ubp3^C469A^ cells were grown in high glucose and the mRNA levels of PFK1, TDH2 and TDH3 were analysed by RT-qPCR. The fold changes in mRNA levels are shown. Data represent mean ± SD. D) Steady-state pyruvate levels in WT and *ubp3Δ* in high glucose. Data represent mean ± SD from three biological replicates (n=3). Also see Appendix Table S3. E) Schematic showing the experimental design for measuring ^13^C label incorporation into glycolytic intermediates and trehalose using a ^13^C_6_ glucose-pulse. F) Changes in ^13^C label incorporation into glycolytic intermediates and trehalose with time, and linearity of label incorporation after a pulse of 1% ^13^C_6_ glucose is shown.

**Figure 2-figure supplement 2:**
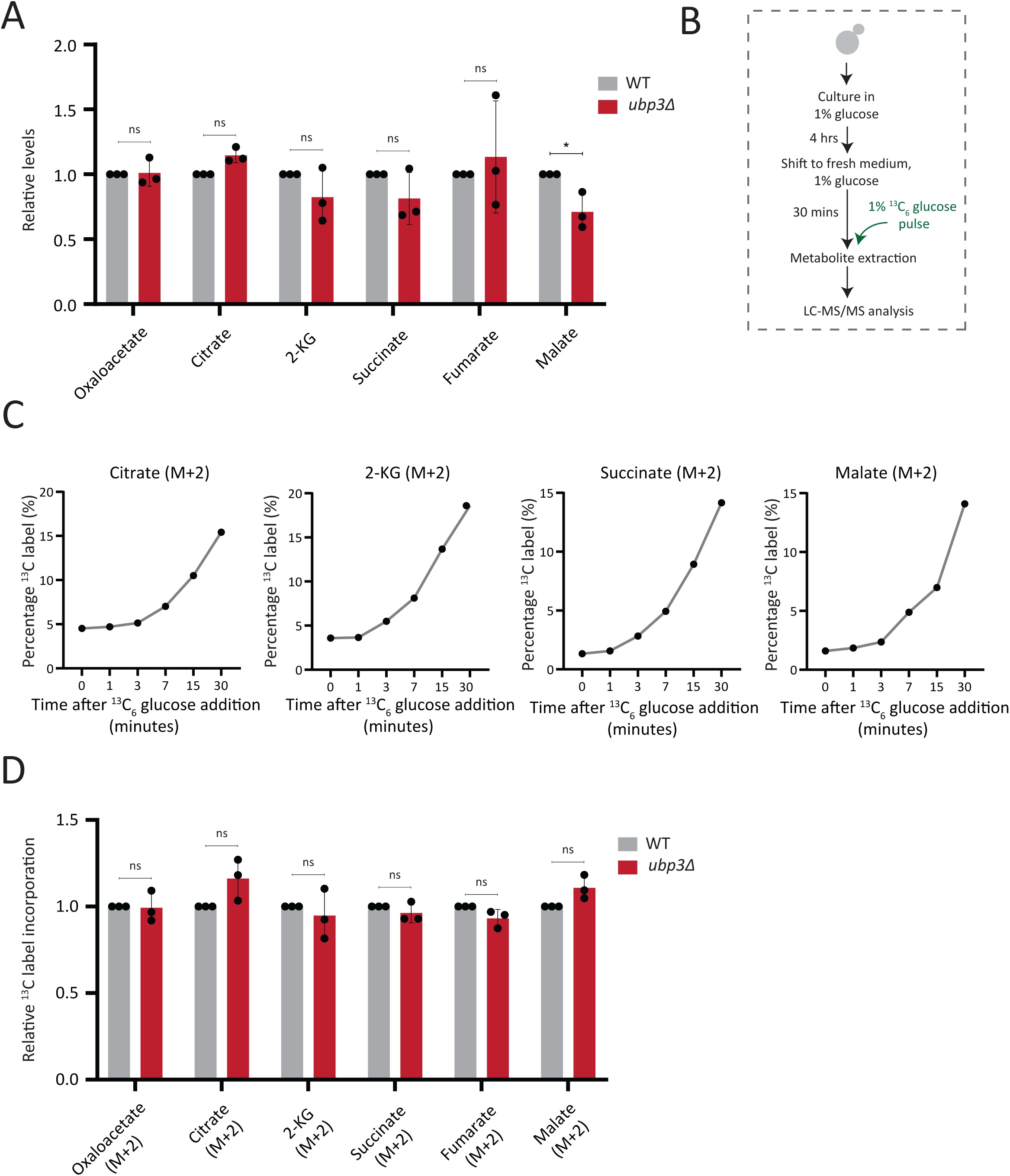
Estimation of steady state levels and flux of TCA cycle intermediates. A) Steady-state TCA metabolite amounts in WT and *ubp3Δ* in high glucose. Relative steady-state levels of TCA cycle intermediates were estimated in WT and *ubp3Δ*. Data represent mean ± SD from three biological replicates (n=3). Also see Appendix Table S3. B) Schematic showing the experimental design for measuring ^13^C label incorporation into TCA cycle intermediates using a ^13^C_6_ glucose-pulse. C) Changes in ^13^C label incorporation into TCA cycle intermediates with time, and linearity of label incorporation after a pulse of 1% ^13^C_6_ glucose is shown. D) Relative TCA cycle flux in WT and *ubp3Δ*. Relative ^13^C-label incorporation into TCA cycle intermediates, after a pulse of 1% ^13^C_6_ glucose is shown. Data represent mean ± SD from three biological replicates (n=3). Also see Appendix Table S3. Data information: *p<0.05

**Figure 3-figure supplement 1:**
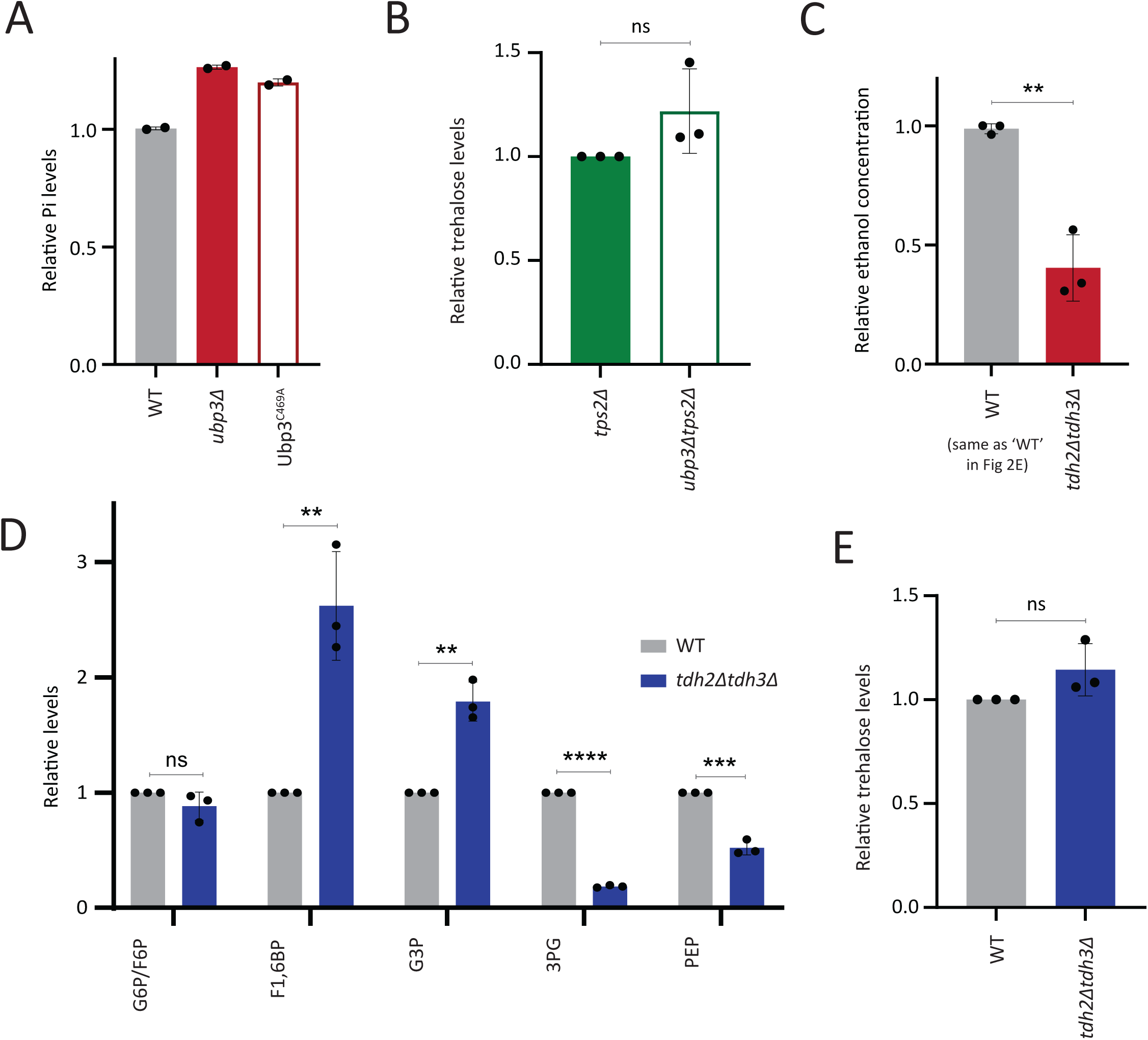
Comparisons of phosphate, ethanol and other metabolites in WT, *ubp3Δ* and GAPDH mutants. A) Loss of Ubp3 deubiquitinase activity and Pi levels. WT, *ubp3Δ,* and Ubp3^C469A^ cells were grown in high glucose and the total free phosphate (Pi) levels were estimated. Data from two independent experiments (n=2) is shown. Data represent mean ± SD. B) Trehalose levels in WT and *ubp3Δ* cells in the absence of Tps2. *tps2Δ* and *ubp3Δtps2Δ* cells were grown in high glucose, and the trehalose levels were estimated using targeted LC-MS/MS. Data represent mean ± SD from three biological replicates (n=3). C) Loss of GAPDH isozymes Tdh2 and Tdh3 and ethanol production. WT and *tdh2Δtdh3Δ* cells were grown in high glucose and the ethanol concentration in the medium was measured. Data are represented as mean ± SD from three biological replicates (n=3). Note: The WT shown here is same as the WT in Figure 2E, the ethanol assays in Figure 2E and Figure S3C were done together and a common WT control was used. D) Steady state levels of glycolytic intermediates: *tdh2Δtdh3Δ* cells have unaltered steady state levels of G6P/F6P, increased levels of F1,6 BP and G3P and decreased levels of 3PG and PEP. WT and *tdh2Δtdh3Δ* cells were grown in high glucose and the steady state amounts of glycolytic intermediates were estimated using targeted LC-MS/MS. Data represent mean ± SD from three biological replicates (n=3). Also see Appendix Table S3. E) Trehalose levels in *tdh2Δtdh3Δ* cells. WT and *tdh2Δtdh3Δ* cells were grown in high glucose and trehalose amounts were estimated using targeted LC-MS/MS. Data represent mean ± SD from three biological replicates (n=3). Data information: *p<0.05, **p<0.01.

**Figure 4-figure supplement 1:**
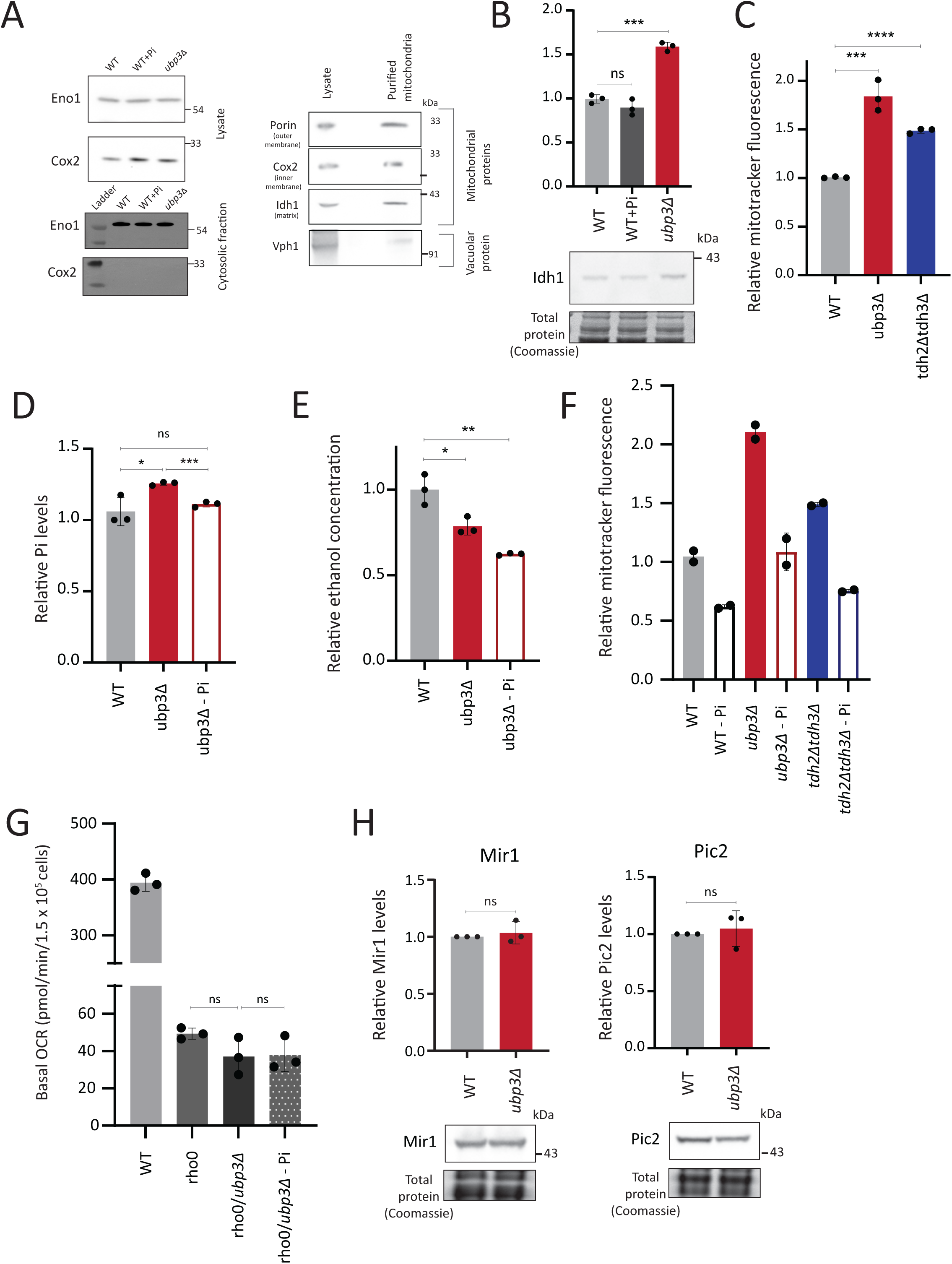
Mitochondrial Pi estimation characterizations and correlations of mitochondrial activity with Pi availability. A) Isolation of cytosolic fraction by centrifugation and lack of vacuolar contamination in mitochondria isolated by immunoprecipitation. Cytosolic fraction was isolated from WT cells grown in high glucose with or without excess Pi and *ubp3Δ* cells. Eno1 was used as a cytosolic marker and Cox2 as mitochondrial marker to ensure purity of the cytosolic fractions. Mitochondria were isolated from WT cells grown in high glucose and the protein levels of Tom20, Cox2, Idh1 and Vph1 and were measured in cell lysate and immunoprecipitated mitochondria by western blot. B) Idh1 levels in *ubp3Δ* cells and WT cells grown in high Pi. WT cells were grown in high glucose, or high glucose and high Pi medium (10mM Pi) and *ubp3Δ* cells were grown in high glucose, and the Idh1 protein levels were estimated by western blot. A representative blot (out of three biological replicates, n=3) is shown in the lower panel. Quantification is shown in the upper panel. Data represent mean ± SD. C) Mitochondrial membrane potential in *tdh2Δtdh3Δ* cells. WT, *ubp3Δ,* and *tdh2Δtdh3Δ* cells were grown inhigh glucose and the mitochondrial membrane potential was measured. Data represent mean ± SD from three biological replicates (n=3). D) Pi levels in *ubp3Δ* cells grown in low Pi (1mM Pi) vs WT cells. WT cells were grown in high glucose and *ubp3Δ* cells were grown in high glucose or high glucose-low Pi, and the total Pi levels were estimated. Data represent mean ± SD from three biological replicates (n=3). E) Ethanol production in *ubp3Δ* cells grown in a low Pi medium. WT cells were grown in high glucose and *ubp3Δ* cells were grown in high glucose or high glucose-low Pi, and the ethanol concentration in the medium was measured. Data represent mean ± SD from three biological replicates (n=3). F) Pi amounts and mitochondrial membrane potential in WT, *ubp3Δ,* and *tdh2Δtdh3Δ* cells. The cells were grown in high glucose or high glucose-low Pi, and the mitochondrial membrane potential was measured. Data from two independent experiments (n=2) is shown. Data represent mean ± SD. G) The Pi mediated change in basal OCR in *ubp3Δ* is dependent on mitochondrial respiration. WT and Rho0 cells were grown in high glucose, Rho0/*ubp3Δ* were grown in high glucose and low Pi, and basal OCR was measured from three independent experiments (n=3). Data represent mean ± SD. H) The protein levels of mitochondrial Pi transporters Mir1 and Pic2 in *ubp3Δ* cells. WT and *ubp3Δ* cells containing endogenously tagged Mir1 and Pic2 at their C terminus with a 6x HA epitope tag were grown in high glucose and Mir1 and Pic2 proteins were measured by western blot. A representative blot (out of three biological replicates, n=3) is shown in the lower panel. Quantification is shown in the upper panel. Data represent mean ± SD. Data information: *p<0.05, **p<0.01, ***p<0.001.

**Figure 5-figure supplement 1:**
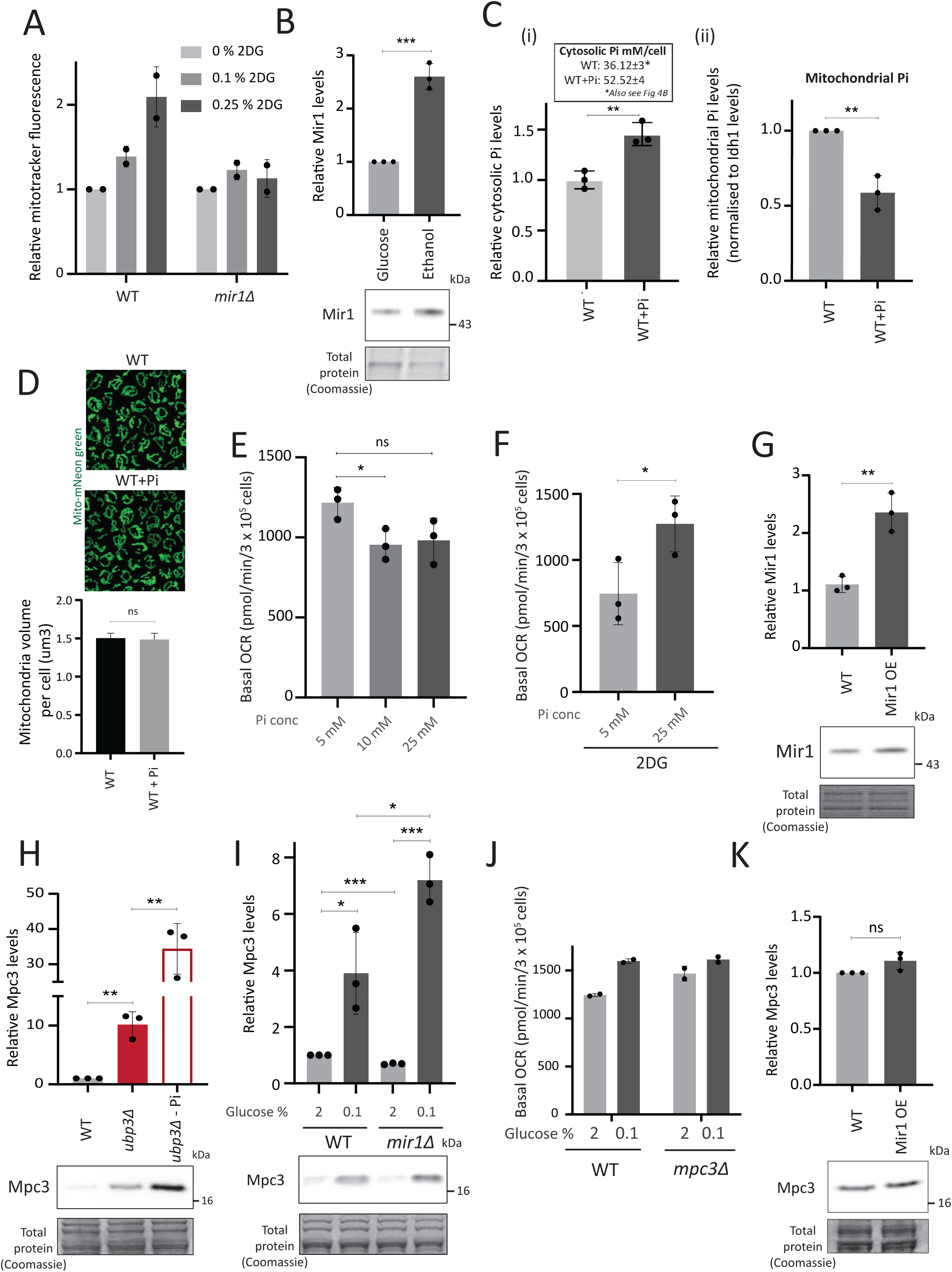
Mitochondrial phosphate and pyruvate transport relationships with mitochondrial activity. A) Requirement of Mir1 for switching to increased mitochondrial activity upon 2-deoxyglucose (2-DG) treatment. WT and *mir1Δ* cells were grown inhigh glucose and treated with 0.1% and 0.25% 2-DG for one hour. The mitochondrial membrane potential was measured from two independent experiments (n=2). Data represent mean ± SD. B) Mir1 protein levels and glucose repression. Cells containing Mir1 endogenously tagged at the C terminus with a 6x HA epitope tag were grown in high glucose or a respiratory medium (2% ethanol). Mir1 protein was estimated by western blot. A representative blot (out of three biological replicates, n=3) is shown in the lower panel. Quantification is shown in the upper panel. Data represent mean ± SD. C) Effect of supplementing Pi in the medium on cytosolic and mitochondrial Pi. WT cells were grown in high glucose, and high glucose-high Pi medium (10 mM Pi). The cytosolic fraction was isolated by centrifugation (see SI appendix), and in separate experiments, mitochondria were isolated by immunoprecipitation from WT and *ubp3Δ* and mitochondrial Pi estimated. (i) Cytosolic Pi levels (relative and absolute) and (ii) mitochondrial Pi levels normalised to Idh1 protein levels is shown. Data represent mean ± SD from three biological replicates (n=3). Also see Figure 4-figure supplement 1A, B. D) Effect of supplementing Pi on the total mitochondrial volume. WT cells with mitochondria targeted mNeon green were grown in high glucose, and high glucose, high Pi medium (10 mM Pi), cells were imaged and the total mitochondrial volume per cell was calculated. E) Increasing Pi concentration in a high glucose medium results in a decrease in basal OCR. WT cells were grown in high glucose medium (YPD-Pi medium), supplemented with 5 mM Pi and at OD_600_∼0.6, the medium was supplemented with Pi (10 mM and 25 mM final concentrations). The basal OCR was measured one hour after Pi supplementation. Data represent mean ± SD from three biological replicates (n=3). E) Effect of increasing the Pi concentration in a high glucose medium, in the presence of 2DG, on basal OCR. WT cells were grown in high glucose medium (YPD-Pi medium), supplemented with 5 mM Pi and at OD_600_∼0.6, the medium was supplemented with Pi (25 mM final concentration) and 2DG (0.25%) for one hour and the basal OCR was measured. Data represent mean ± SD from three biological replicates (n=3). F) Overexpression of Mir1. C terminal 6x HA epitope-tagged Mir1 was expressed under G6PD promoter in cells with Mir1 endogenously tagged at the C terminus with a 6x-HA epitope tag. WT cells (carrying an empty vector, and expressing Mir1-HA under the endogenous Mir1 promoter) and Mir1-OE cells were grown in high glucose. The protein levels of Mir1 were estimated by western blot, and clones with an ∼2-fold increase in Mir1 were selected. A representative blot (out of three biological replicates, n=3) is shown in the lower panel. Quantification is shown in the upper panel. Data represent mean ± SD. G) Mpc3 protein levels in *ubp3Δ* cells. WT and *ubp3Δ* cells containing endogenously tagged Mpc3 at the C terminus with a 3x FLAG epitope tag, were cultured in high glucose (2% glucose) and high glucose-low Pi (2% glucose, 1 mM Pi). The Mpc3 protein levels were measured by western blot. A representative blot (out of three biological replicates, n=3) is shown in the lower panel, quantifications shown in the upper panel. Data represent mean ± SD. H) Mpc3 protein amounts upon shifting WT and *mir1Δ* cells to a low-glucose medium. WT and *mir1Δ* cells containing Mpc3-FLAG, were cultured in high glucose (2% glucose) and shifted to low (0.1%) glucose for 1 hour. Mpc3 protein was measured by western blot. A representative blot (out of three biological replicates, n=3) is shown in the lower panel, quantifications shown in the upper panel. Data represent mean ± SD. I) Requirement of Mpc3 for the increase in basal OCR upon shifting to low glucose. WT and *mpc3Δ* cells were cultured in high glucose (2% glucose) and shifted to low (0.1%) glucose for 1 hour. The basal OCR from two independent experiments (n=2), normalized to the OD600 is shown. Data represent mean ± SD. J) Mpc3 levels and Mir1 overexpression. WT cells and Mir1-OE cells containing endogenously tagged Mpc3 ate the C terminus with a 3x FLAG tag were grown in high glucose. The protein levels of Mpc3 were estimated by western blot. A representative blot (out of three biological replicates, n=3) is shown in the panel. Quantification is shown in the upper panel. Data represent mean ± SD. Data information: *p<0.05, **p<0.01, ***p<0.001.

