## Supplementary material for "The deubiquitinase Ubp3/Usp10 constrains glucose-mediated mitochondrial repression via phosphate budgeting": SI Appendix

1. Institute for Stem Cell Science and Regenerative Medicine (DBT-inStem)
   GKVK Post Bellary Road
   Bangalore 560065
2. Manipal Academy of Higher Education, Manipal, Karnataka 576104, India

**<u>Table of contents</u>**

**Extended methods - Page 2**

**Appendix Table S1 - Page 9**

**Appendix Table S2 - Page 11**

**Appendix Table S3 - Page 11**

**SI references - Page 12**

### Extended methods

#### Media and growth conditions

Media used in this study are high glucose (1% yeast extract, 2% peptone, and 2% glucose), low glucose (1% yeast extract, 2% peptone, and 0.1% glucose), and ethanol (1% yeast extract, 2% peptone, and 2% ethanol). The YPD-low Pi medium was prepared as previously described (Kaneko et al., 1982). Briefly, 1 L YP medium was prepared with 10g yeast extract and 20g peptone in 800ml water. To this, 10 ml of 1M $MgS0_4$ and 10ml aqueous ammonia were added and incubated for 30 mins at room temperature (RT), to precipitate inorganic phosphate. The precipitate was filtered and the pH of the clear solution adjusted to 5.8 using HCl, and the volume made up to 1 L. This medium was autoclaved and sterile dextrose was added to prepare the YPD-no Pi medium. The low Pi medium was prepared by adding an indicated concentration of filter-sterilized $KH_2PO_4$ solution. For experiments involving shift to low glucose, cells were grown in high glucose (2%) and at $OD_{600}$ ~0.6, cells were pelleted and shifted to low glucose (0.1%), for one hour. For experiments involving a shift to a no-Pi or low glucose medium, cells were subcultured in high glucose till $OD_{600}$~0.6, pelleted by centrifugation (1000 x g, 2 min at RT), washed, and shifted to no-Pi or low glucose medium for 1 hour. For experiments involving measurement of basal OCR at different Pi concentrations (Figures S5D, S5E), cells were grown in 5 mM Pi medium (prepared by removing Pi from YPD medium and adding $KH_2PO_4$ at the required concentration) and at $OD_{600}$ ~0.6, the cells were supplemented with additional Pi at required final concentrations. MG132 was used at a final concentration of 100 µM and the experiments involving MG132 addition were performed in pdr5Δ background to prevent MG132 efflux by Pdr5 transporter. Experiments involving heat stress were performed by incubating cells at 42°C for one hour.

#### CRISPR-Cas9 based mutagenesis

To generate the $Ubp3^{C469A}$ strain, cells were transformed a constitutive Cas9 expressing plasmid (Addgene plasmid 43802, (Dicarlo et al., 2013)). Guide RNA (gRNA) sequences (forward and reverse) were annealed and cloned into a pMEL13 plasmid. Strains were then transformed with gRNA plasmid and homology repair (HR) fragments with the base pair mutation. The HR fragment was generated by site-directed mutagenesis PCR. Positive clones were selected using drug, and sequenced to confirm the mutation. Oligonucleotides used are listed in appendix table S2.

**Sample preparation for mitotracker fluorescence assay**

Cells were grown to $OD_{600}$~0.6, 1 $OD_{600}$ cells were collected by centrifugation (1000 x g, 1 min at RT), and resuspended in 1 ml fresh media. Mitotracker CMXRos (M7512 ThermoFisher) was added to a final concentration of 200 nM and incubated at 30°C in a shaking incubator. Cells were washed and resuspended in fresh media and fixed with 2% formaldehyde for 20 minutes. Fixed cells were washed in 1xPBS, pH 7.4, and resuspended in 1 ml 1xPBS. 300 µl of this was aliquoted onto 96 well plates in replicates.

**Mitotracker based screen to identify DUBs that regulate mitochondrial repression**

The deubiquitinase deletions were made by PCR mediated gene deletion, DUB deletion mutants generated were grown in high glucose and the mitochondrial membrane potential measured by the mitotracker assay described earlier. For performing the screen, the 19 DUB knockouts were divided into two groups- each containing 10 and 9 DUB KOs respectively and the screen was performed separately for each group in replicates. WT cells grown in high glucose indicated the basal mitochondrial potential and WT cells grown in ethanol was used as a positive control for each group. The mitotracker fluorescence intensity were normalised to $OD_{600}$ of each mutant and the fluorescence intensity relative to the WT cells were calculated. The relative fluorescence intensity for the replicate samples are shown in supplementary figure S1A, mean fluorescence intensities of the replicates (relative to WT) were calculated and plotted as heat maps (Figure 1C).

**Serial dilution growth assay**

For serial dilution-based growth assays, cells were grown in standard high glucose to $OD_{600}$~0.8, collected by centrifugation (1000 x g, 1 min at RT), washed with water, serial dilutions made

($OD_{600}$=1, 0.1, 0.01, 0.001, 0.0001), 5 µl of each dilution were spotted onto agar plates, incubated at 30°C and growth monitored.

**Protein extraction and western blotting**

The cells were grown in the indicated medium to $OD_{600}$~0.8 and pelleted by centrifugation (1000 x g, 2 min at RT). Total protein was precipitated and extracted using trichloroacetic acid (TCA), and resuspended in SDS/glycerol buffer. The supernatant was collected after centrifugation, and total proteins were estimated by BCA assay (BCA assay kit, G-Biosciences). Protein samples were normalized to ensure the same protein amounts in all the samples, in SDS/glycerol buffer. Samples were resolved on 4–12% bis-tris gels (Invitrogen, NP0336BOX), using MOPS running buffer (50 mM tris, 50 mM MOPS, 0.1% SDS, and 1 mM EDTA), the gels cut so that the relevant portion of the gel is transferred to a nitrocellulose membrane (GE Healthcare, 10600003), while a lower or higher region was stained with coomassie blue for protein normalization, and blots were developed using the following antibodies: anti-HA mouse (Sigma-Aldrich 11583816001), anti-FLAG mouse (Sigma-Aldrich F1804), anti-Cox2 mouse (Invitrogen MTCO2 459150), anti-ubiquitin (P4D1 mouse mAb, CST), anti-Idh1 goat (Sigma-Aldrich SAB2501682). Horseradish peroxidase-conjugated secondary antibody was from Sigma-Aldrich (mouse and rabbit) and Thomas scientific (goat). Chemiluminescence was detected by using Western Bright ECL HRP substrate, Advansta, K12045.

**Seahorse assay**

1 ml of the XF Calibrant solution was aliquoted into each well of the utility plate and the sensor cartridge plate was hydrated overnight as per the manufacturer's instructions. The sensor cartridge was loaded with complex IV inhibitor sodium azide and equilibrated in the Seahorse XFe24 analyzer one hour prior to the start of the assay. The culture plate was coated with 50 µl poly-L-lysine and incubated for 1 hour at RT. Excess poly-L-lysine was removed and the plate dried at 30°C for 30 minutes. The cells were grown in standard high glucose to an $OD_{600}$~0.6. The samples were aliquoted to poly-L-lysine coated wells at a final cell number in each well of ~3x10^5. The plate was centrifuged for 2 min at 100xg (acceleration 2, brake 2) and incubated at 30°C for 30 minutes. The plate was loaded to the Seahorse XFe24 analyzer and basal OCR measured over time. A minimum of three measurements were taken with intermittent 2 min

mixing and waiting steps. This was followed by sodium azide injection to the sample wells, and three measurements were taken with intermittent 2 min mixing and waiting steps.

**Sample preparation for ATP estimation assay**

Cells were grown in high glucose and at $OD_{600}$ ~0.8, 10 $OD_{600}$ were pelleted at 4°C. The pellet was treated with 300 µl ice-cold TCA (5%), resuspended and incubated in ice for 15 mins. The suspension was diluted such that the final TCA concentration is 0.1% using 20 mM Tris-HCl, Ph 7.0. For calculating the contribution of mitochondria to total ATP, cells were grown in high glucose and at $OD_{600}$ ~0.6, sodium azide was added at a final concentration of 1 mM and incubated for 45 minutes at 30°C in a shaking incubator. The reaction mixture for ATP measurements were prepared as per manufacturer's instructions and luminescence was measured using Sirius luminometer (Tiertek Berthold). The relative ATP concentrations were calculated, graphs were plotted and using GraphPad prism 9.0.1.

**Sample preparation for ethanol estimation**

Cells were grown in appropriate media conditions to an $OD_{600}$~1, 10 ml of cells collected, and centrifuged at 3000 x g at 4°C. 5 ml of the supernatant was collected in a fresh tube and 1ml of Tri-n-butyl phosphate (TBP) was added. The mixture was vortexed for 5 minutes and centrifuged at 3420xg for 5 minutes to separate the phases. 500 µl of the top, clear phase was transferred to a fresh tube and 500 µl of potassium dichromate reagent was added (10% w/v of $K_2Cr_2O_7$ in 5 M of $H_2SO_4$), followed by vortexing for 1 minute. The mixture was incubated at RT for 10 minutes and 200 µl of the bottom layer was transferred to a 96 well plate. The absorbance in each well was measured at 595 nm. A standard curve was plotted using medium containing different concentrations of added ethanol, and was used to calculate the ethanol concentration in samples. For calculating the rate of ethanol production, WT and *ubp3Δ* cells were grown in high glucose and at $OD_{600}$~0.6, equal number of cells were shifted to fresh high glucose medium. The supernatant from 10ml cells were collected, and ethanol concentration in the medium was estimated at different time points after the shift (normalised to $OD_{600}$ at each time point, to consider the growth difference between WT and *ubp3Δ* cells).

**Sample preparation for Pi estimation by malachite green assay**

The cells were grown to $OD_{600}$~0.8, pelleted by centrifugation (1000 x g, 2 min at RT), pelleted cells were washed and resuspended in 1 ml ice-cold HPLC grade water (to avoid Pi contamination) and lysed by bead beating (3 x 20s, 1 min intermittent cooling). Lysates were centrifuged (14000 rpm, 1 min at 4°C) and 800 µl of the clear supernatant collected. Protein concentrations of the supernatant were estimated using BCA assay (BCA assay kit, GBiosciences). Lysates corresponding to 1µg total protein was used for the Pi estimation assay.

**Metabolite extraction and analysis by LC-MS/MS:**

**Steady state measurements of glycolytic intermediates**

For measuring steady-state levels of glycolytic intermediates, trehalose, and PPP intermediates, cells were grown in standard high glucose medium to an $OD_{600}$~0.8, 5 $OD_{600}$ cells were collected, quenched in 60% methanol at -40°C, and metabolites were extracted as described in (Walvekar et al., 2018). The extracted metabolites were separated by Synergi 4µm Fusion-RP 80 Å LC column (150 x 4.6 mm, Phenomenex) on Waters Acquity UPLC system, and measured as described earlier (Walvekar et al., 2018), in negative polarity mode. Solvents used: 5 mM ammonium acetate in water (solvent A) and 100% acetonitrile (solvent B). ABSciex QTRAP 6500 mass spectrometer was used and data acquisition was done using Analyst 1.6.2 (Sciex). Detailed flow parameters are described elsewhere (Walvekar et al., 2018). MultiQuant version 3.0.1 was used for data analysis. The parent and product ion masses used for the analysis are listed in appendix table S3. The peak areas for individual metabolites were calculated and values were plotted relative to WT. The retention time for different metabolites and peak area obtained after analysis are listed in supplement file 1. The statistical significance was calculated using unpaired Student's t-test (GraphPad prism 9.0.1).

**Measurement of $^{13}C$ carbon flux into glycolysis and trehalose**

The carbon flux into glycolysis and trehalose was estimated by measuring the relative incorporation of $^{13}C$ label into glycolytic intermediates and trehalose after a pulse of $^{13}C_6$

glucose. Cells were grown in high glucose medium with 1% glucose and at $OD_{600}$~0.6, the cells were shifted to fresh medium with 1% glucose. The $OD_{600}$ was measured 30 minutes after the shift and cells corresponding to ~5 $OD_{600}$ were collected, and $^{13}C_6$ glucose (Cambridge Isotope Laboratories, CLM-1396) was pulsed at a final concentration of 1%, making the total glucose concentration in the medium 2% (1% labelled + 1% unlabelled). Note: Since label saturation into glycolytic intermediates happens instantaneously (within seconds) after $^{13}C$ glucose pulse, the time points for sample collection should be determined prior to the experiment by plotting total label percentage over time for individual metabolites to calculate the time point at which label saturation takes place. For measuring $^{13}C$ label incorporation, the cells were collected at 3s (for F1,6BP, G3P, 3PG and PEP), 10s (for G6P/F6P) and 4 mins (for trehalose) after $^{13}C_6$ glucose pulse, based on the percentage label incorporation plotted in figure S2C. The cells were quenched in 60% methanol at -40°C, and metabolites were extracted as described in (Walvekar et al., 2018). The extracted metabolites were detected as described earlier. The relative $^{13}C$ label incorporation in *ubp3Δ* cells were measured by calculating the area under the peak for individual $^{13}C$ labelled metabolites (using MultiQuant version 3.0.1) and dividing it by the peak area of that specific metabolite in the WT sample. The change in $^{13}C$ labelled metabolites relative to WT samples were plotted. For calculating the total $^{13}C$ label percentage of a metabolite, the peak area of $^{13}C$ labelled metabolites were divided by the sum of peak areas of that metabolite (labelled plus unlabelled) and expressed as percentage. The total label percentage was plotted over time for individual metabolites to calculate the time point at which label saturation takes place (Figure S2C). The retention time for different labelled and unlabelled metabolites and peak area obtained after analysis are listed in supplement file 1. The statistical significance was calculated using unpaired Student's t-test (GraphPad prism 9.0.1).

### Sample preparation for cytosolic fraction separation and mitochondria isolation by immunoprecipitation

Cells with Tom20 (for mitochondria isolation) and Eno1 (for cytosolic fraction isolation) endogenously tagged at the C terminus with a 3X FLAG epitope tag were grown in 300 ml high glucose medium and at $OD_{600}$~ 0.8, the cells were pelleted at 1500 x g for 5 minutes at 4°C. The cells were washed in water, pelleted and weighed. The cells were then incubated in Tris-DTT buffer (0.1 M Tris-$SO_4$, 10 mM DTT, pH 9.4) (5 ml/g pellet weight) for 15 mins at 30°C in a shaking incubator. The cells were washed in SEH buffer (0.6 M sorbitol, 20 mM HEPES-KOH, 2mM $MgCl_2$, pH 7.4) (5 ml/g pellet weight) and incubated with Zymolyase 20T (0832092, MP biomedicals) dissolved in SEH buffer for 60 minutes, at 30°C in a shaking incubator. The spheroplasts produced by Zymolyase treatment were collected by centrifugation at 4500 x g for 5 mins at 4°C, washed with ice-cold SEH buffer (5 ml/g pellet weight) and resuspended in ice-cold SEH buffer with protease inhibitors (1 mM PMSF, 0.5 mg/ml pepstatin A and 0.5 µg/ml leupeptin) (5 ml/g pellet weight). The resuspended spheroplasts were homogenised using a Dounce homogeniser, centrifuged at 1500 x g for 5 mins and the supernatant was collected. The supernatant was centrifuged at 12000 x g for 10 mins at 4°C. The supernatant cytosolic fraction was collected and the pellets were resuspended in 700 µl ice-cold SEH buffer to obtain the mitochondria-enriched fraction. To purify mitochondria by immunoprecipitation, the mitochondria-enriched fraction was incubated with 1.5 mg Dynabeads protein G (Invitrogen) conjugated with 10 µg of anti-FLAG antibody (mouse, Sigma-Aldrich F1804), for 60 minutes at 4°C. The beads were separated using a magnet, washed 3 times in ice-cold SEH buffer and the mitochondria were eluted in 200 ul SEH buffer with FLAG peptide (1.5 mg/ml).

**Mitotracker assay using isolated mitochondria**

For mitotracker assays with isolated mitochondria (Figure 5D), the mitochondria enriched fraction resuspended in SEH buffer was centrifuged twice- 700 x g for 5mins and 1500 x g for 5 mins at 4°C. The supernatant was collected and centrifuged at 12000 x g for 10 mins at 4°C and the pellet was resuspended in ice-cold SEH buffer. The protein concentration of this crude mitochondria was estimated by BCA protein estimation assay and mitochondria corresponding to 5 µg protein was incubated in 200 µl SEH buffer containing 1 mM pyruvate, 1 mM malate, 0.5 mM ADP and 0 – 50 mM $KH_2PO_4$ for 30 minutes at 30°C. Mitotracker CMXROS was added at a final concentration of 200 nM, followed by incubation at 30°C for 30 minutes. The fluorescence intensities were measured using Thermo Varioscan™ LUX multimode plate reader at 579/599 excitation/emission, and fluorescence intensities relative to samples containing 0 mM KH2PO4 was calculated. Statistical significance was calculated using unpaired Student's t-test (GraphPad prism 9.0.1).

**Appendix Table S1: List of strains used in this study**

| SL no. | Strain name | Genotype | Description | Source |
|---|---|---|---|---|
| 1 | CEN.PK a (WT) | Mat a | haploid strain of CEN.PK MAT 'a' mating type | (J.P. van Dijken et al., 2000) |
| 2 | DUBs KO strains | CEN.PK a::hphMX6 | Deletion of the individual deubiquitinases (see supplement figure S1) | This study |
| 3 | Ubp3 $_{C469A}$ | CEN.PK a UBP3 $^{C469A}$ | Catalytically inactive Ubp3 mutant | This study |
| 4 | *Δatp1* | CEN.PK a *Δatp1*::natMX6 | Deletion of ATP1 gene | This study |
| 5 | *Δatp1 Δubp3* | CEN.PK a *Δubp3::hphMX6 Δatp1:: natMX6* | Deletion of ATP1 gene in a UBP3 deletion strain | This study |
| 6 | *Δatp10* | CEN.PK a *Δatp10*::natMX6 | Deletion of ATP10 gene | This study |
| 7 | *Δatp10 Δubp3* | CEN.PK a *Δubp3::hphMX6 Δatp10:: natMX6* | Deletion of ATP10 gene in a UBP3 deletion strain | This study |
| 8 | Pfk1-FLAG | CEN.PK a *PFK1 3xFLAG::natMX6* | C terminal tagged Pfk1 | This study |
| 9 | Pfk1-FLAG *Δubp3* | CEN.PK a *PFK1-3xFLAG::natMX6 Δubp3::hphMX6* | C terminal tagged Pfk1 in a UBP3 deletion strain | This study |
| 10 | Tdh2-FLAG | CEN.PK a *TDH2-3xFLAG::natMX6* | C terminal tagged Tdh2 | This study |
| 11 | Tdh2-FLAG *Δubp3* | CEN.PK a *TDH2-3xFLAG::natMX6 Δubp3::hphMX6* | C terminal tagged Tdh2 in a UBP3 deletion strain | This study |
| 12 | Tdh3-FLAG | CEN.PK a *TDH3-3xFLAG::natMX6* | C terminal tagged Tdh3 | This study |
| 13 | Tdh3-FLAG *Δubp3* | CEN.PK a *TDH3-3xFLAG::natMX6 Δubp3::hphMX6* | C terminal tagged Tdh3 in a UBP3 deletion strain | This study |
| 14 | Eno1-FLAG | CEN.PK a *ENO1-3xFLAG::natMX6* | C terminal tagged Eno1 | This study |

| | | | | |
|---|---|---|---|---|
| 15 | Eno1-FLAG *Δubp3* | CEN.PK a *ENO1-3xFLAG::natMX6 Δubp3::hphMX6* | C terminal tagged Eno1 in a UBP3 deletion strain | This study |
| 16 | Eno2-FLAG | CEN.PK a *ENO2-3xFLAG::natMX6* | C terminal tagged Eno2 | This study |
| 17 | Eno2-FLAG *Δubp3* | CEN.PK a *ENO2-3xFLAG::natMX6 Δubp3::hphMX6* | C terminal tagged Eno2 in a UBP3 deletion strain | This study |
| 18 | *Δtdh2 Δtdh3* | CEN.PK a *Δtdh2::natMX6 Δtdh3:: kanMX6* | Deletion of TDH2 gene in a TDH3 deletion strain | This study |
| 19 | *Δtps2* | CEN.PK a *Δtps2::hphMX6* | Deletion of TPS2 gene | This study |
| 20 | *Δtps2 Δubp3* | CEN.PK a *Δtps2::hphMX6 Δubp3::natMX6* | Deletion of UBP3 gene in a TPS2 deletion strain | This study |
| 21 | Mir1-HA | CEN.PK a *MIR1-6xHA::natMX6* | C terminal tagged Mir1 | This study |
| 22 | Mir1-HA *Δubp3* | CEN.PK a *MIR1-6xHA::natMX6 Δubp3::hphMX6* | C terminal tagged Mir1 in a UBP3 deletion strain | This study |
| 23 | Pic2-HA | CEN.PK a *PIC2-6xHA::natMX6* | C terminal tagged Pic2 | This study |
| 24 | Pic2-HA *Δubp3* | CEN.PK a *PIC2-6xHA::natMX6 Δubp3::hphMX6* | C terminal tagged Pic2 in a UBP3 deletion strain | This study |
| 25 | *Δmir1* | CEN.PK a *Δmir1::natMX6* | Deletion of Mir1 | This study |
| 26 | *Δmir1 Δubp3* | CEN.PK a *Δmir1::natMX6 Δubp3::hphMX6* | Deletion of Mir1 in a UBP3 deletion strain | This study |
| 27 | WT+ Empty vector | CEN.PK a *pG6PD:: kanMX6* | haploid strain of CEN.PK MAT 'a' with an empty vector with kanamycin resistance | This study |
| 28 | Mir1-HA OE | CEN.PK a *pG6PD-Mir1-6xHA:: kanMX6* | Mir1-HA overexpression under the constitutive G6PD promoter | This study |

| | | | | |
|---|---|---|---|---|
| 29 | Mpc3-FLAG | CEN.PK a *MPC3-3xFLAG::natMX6* | C terminal tagged Mpc3 | This study |
| 30 | Mpc3-FLAG *Δubp3* | CEN.PK a *MPC3-3xFLAG::natMX6 Δubp3::hphMX6* | C terminal tagged Mpc3 in a UBP3 deletion strain | This study |
| 31 | Mpc3-FLAG Mir1-HA OE | CEN.PK a *MPC3-3xFLAG::natMX6 pG6PD-Mir1-6xHA:: kanMX6* | C terminal tagged Mpc3 in Mir1-HA overexpression | This study |
| 32 | *Δmpc3* | CEN.PK a *Δmpc3::natMX6* | Deletion of Mpc3 | This study |
| 33 | Tom20-FLAG | CEN.PK a Tom20-*3xFLAG::natMX6* | C terminal tagged Tom20 | This study |
| 34 | Tom20-FLAG Vph1-HA | CEN.PK a Tom20-*3xFLAG::natMX6 Vph1-6xHA:: hphMX6* | C terminal tagged Vph1 in c terminal tagged Tom20 | This study |
| 35 | Mito-Mneon green | CEN.PK a *HO::$P_{CYC1}$-SU9mNeongreen-$T_{CYC1}$-KanMX6* | Mneon gene with a mitochondria targeted sequence at the N terminus | (Dua et al., 2022) |
| 36 | cox2-62 | *leu2 Δarg8 ΔURA3 ura3-52 kar1-1 ade2-101* | cox2-62 ρ+, Cox2 with deletion of -295 to +363 relative to AUG | (Bonnefoy et al., 2001) |
| 37 | W303 | MAT a *leu2-3,-112;his3-11,-15;trp11;ura3-1;ade2-1;can1-100* | WT W303 strain | (Ralser et al., 2012) |
| 38 | W303 *Δubp3* | MAT a *leu2-3,-112;his3-11,-15;trp11;ura3-1;ade2-1;can1-100Δubp3::hphMX6* | Deletion of Ubp3 in W303 | This study |
| 39 | BY4742 | MAT α *his3Δ1:leu2Δ0:lys2Δ0:MET15:ura3Δ0* | WT BY4742 strain | (Winston et al., 1995) |
| 40 | BY4742 *Δubp3* | MAT α *his3Δ1:leu2Δ0:lys2Δ0:MET15:ura3Δ0 Δubp3::hphMX6* | Deletion of Ubp3 in BY4742 | This study |
| 41 | Σ1278 | MAT a | WT Σ1278 strain | Isolate via *Fink lab* |

| | | | | |
|---|---|---|---|---|
| 42 | Σ1278 *Δubp3* | MAT a *ura3-52 Δubp3::hphMX6* | Deletion of Ubp3 in BY4742 | This study |
| 43 | S288C | MAT a | WT S288C strain | This study |
| 44 | Rho0 | CEN.PK Mat a | Lacks mitochondrial DNA, generated using EtBr treatment | This study |
| 45 | Rho0 *Δubp3* | CEN.PK Mat a | Deletion of Ubp3 in Rho0 strain | This study |

**Appendix Table S2: Oligonucleotides used for CRISPR-Cas9 based mutagenesis**

| | |
|---|---|
| Ubp3C469A gRNA forward | AGAACTCATAAAACAAATGTgtttt |
| Ubp3C469A gRNA reverse | ACATTTGTTTTATGAGTTCTgatca |
| HR fragment Ubp3C469A forward | CAAAATACCAGTCCATTCCATTATTCCAAGAGGCATAATTAACAGAGCCAACATTGCTTTTATGAGTTCT |
| HR fragment Ubp3C469A reverse | ACGTTAATTACATCAATAAATGGCTTACAGTAGAGTAACACTTGTAACACAGAACTCATAAAAGCAATGT |

**Appendix Table S3: List of parent ion mass and product ion mass (Q1/Q3) used for detection of metabolites**

| **Metabolite** | **Parent Ion (Q1) mass** | **Product Ion (Q3) mass** | **Collision energy (V)** | **Retention time** |
|---|---|---|---|---|
| Glucose 6-phosphate (G6P) / Fructose 6-phosphate (F6P) | 259 | 97 | -20 | 2.68 |
| 13C_G6P/F6P_6 | 265 | 97 | -20 | 2.68 |
| Fructose 1,6-bisphosphate (F16BP) | 339 | 97 | -20 | 2.42 |

| | | | | |
|---|---|---|---|---|
| 13C_F16BP_6 | 345 | 97 | -20 | 2.42 |
| Trehalose | 341.3 | 179.3 | -17 | 3.85 |
| 13C_Trehalose_6 | 347.3 | 185.3 | -17 | 3.85 |
| 13C_Trehalose_12 | 353.3 | 185.3 | -17 | 3.85 |
| Ribose 5-phosphate (R5P) | 229 | 97 | -20 | 2.69 |
| Sedoheptulose-7-phosphate (S7P) | 289 | 97 | -20 | 2.68 |
| Glyceraldehyde 3-phosphate (G3P) | 169 | 97 | -20 | 2.65 |
| 13C_G3P_3 | 172 | 97 | -20 | 2.65 |
| Phosphoenol pyruvate (PEP) | 167 | 79 | -12 | 2.45 |
| 13C_PEP_3 | 170 | 79 | -12 | 2.45 |
| 3-phosphoglycerate (3PG) | 185 | 97 | -20 | 2.53 |
| 13C_3PG_3 | 188 | 97 | -20 | 2.53 |
| Pyruvate | 299.1 | 91.1 | 28 | 8.89 |
| Citrate | 508 | 385 | 7 | 8.04 |
| 13C_ Citrate _2 | 510 | 387 | 7 | 8.03 |
| 13C_ Citrate _3 | 511 | 388 | 7 | 8.03 |
| 13C_ Citrate _4 | 512 | 389 | 7 | 7.69 |
| 13C_ Citrate _5 | 513 | 390 | 7 | 7.91 |

| | | | | |
|---|---|---|---|---|
| 13C_ Citrate _6 | 514 | 391 | 7 | 8.42 |
| 2-Ketoglutarate (2-KG) | 462 | 339 | 11 | 8.68 |
| 13 C_2-KG_1 | 463 | 340 | 11 | 8.67 |
| 13 C_2-KG_2 | 464 | 341 | 11 | 8.67 |
| 13 C_2-KG_3 | 465 | 342 | 11 | 8.63 |
| 13 C_2-KG_4 | 466 | 343 | 11 | 8.6 |
| 13 C_2-KG_5 | 467 | 344 | 11 | 8.69 |
| Succinate | 329 | 206 | 15 | 8.21 |
| 13C_Succinate_1 | 330 | 207 | 15 | 7.32 |
| 13C_Succinate_2 | 331 | 208 | 15 | 7.32 |
| 13C_Succinate_3 | 332 | 209 | 15 | 6.97 |
| 13C_Succinate_4 | 333 | 210 | 15 | 6.96 |
| Fumarate | 327 | 91.2 | 34 | 7.55 |
| 13C_Fumarate_1 | 328 | 91.2 | 34 | 7.74 |
| 13C_Fumarate_2 | 329 | 91.2 | 34 | 7.32 |
| 13C_Fumarate_3 | 330 | 91.2 | 34 | 7.32 |
| 13C_Fumarate_4 | 331 | 91.2 | 34 | 7.32 |
| Malate | 345 | 91.2 | 33 | 7.09 |
| 13C_Malate_1 | 346 | 91.2 | 33 | 7.08 |
| 13C_Malate_2 | 347 | 91.2 | 33 | 7.08 |
| 13C_Malate_3 | 348 | 91.2 | 33 | 7.2 |

| | | | | |
|---|---|---|---|---|
| 13C_Malate_4 | 349 | 91.2 | 33 | 6.61 |
| Oxaloacetate | 448 | 325 | 10 | 9.92 |
| 13C_Oxaloacetate_1 | 449 | 326 | 10 | 8.67 |
| 13C_Oxaloacetate_2 | 450 | 327 | 10 | 8.67 |
| 13C_Oxaloacetate_3 | 451 | 328 | 10 | 8.63 |
| 13C_Oxaloacetate_4 | 452 | 329 | 10 | 8.6 |
